# Multi-lineage emergence of USA300 MRSA in in Chile and a potential role for environmental copper resistance

**DOI:** 10.64898/2026.10.22.761065

**Authors:** Amy Elizabeth Campbell, José Rodrigo W. Martínez, Lorena Díaz, Colleen Bianco, Erin Theiller, Swetha Rajagopal, Alexander Arvanitis, Alejandra Londoño Jiminez, Ahmed M. Moustafa, Lina Rivas, Patricia García, Cesar A. Arias, José M. Munita, Paul J. Planet, the MICROB-R Consortium

**Author notes:** Authors contributed equally (Amy Elizabeth Campbell, José Rodrigo W. Martínez).

## Abstract

Methicillin-resistant *S. aureus* (MRSA) is prevalent worldwide, but individual strains of MRSA tend to spread and expand in relatively confined geographic areas. The USA300 lineage of MRSA is most well-known for its rapid epidemic spread in North America during the first decade of the 2000’s leading to an overall increase in serious skin infections and the emergence of methicillin resistance in the majority of community acquired *S. aureus* infections. A distinct but closely related South American epidemic clade of MRSA USA300 (USA300-SAE), first reported in 2006, proliferated rapidly in Northern South America but was rarely found in other Latin American countries. In a new survey of 274 invasive MRSA isolates from 2022 collected in Chile, we saw a marked increase in USA300-SAE in Southern South America. We found that Chilean USA300-SAE is phylogenetically diverse and was likely introduced multiple times from multiple different South American countries over the course of more than a decade, but only recently became a highly prevalent clone. Using novel molecular diagnostic methods for isolates collected in 2023, we estimate that USA300-SAE made up more than 22% of invasive MRSA strains in Chile in that year and is likely still increasing. We show that copper-associated genes in the signature copper and mercury resistance mobile element (COMER) of USA300-SAE are strongly conserved, and that isogenic mutants of the copper resistance genes in COMER are highly susceptible to copper killing at concentrations seen in tap water. These findings suggest that environmental exposure to copper may provide a fitness advantage that enhances spread of this lineage.

## INTRODUCTION

Methicillin-resistant *Staphylococcus aureus* (MRSA) has been a major global health concern since its emergence in the mid-20^th^ century, becoming endemic in many regions of the world. Though resistance to most beta-lactam antibiotics is the phenotypic hallmark of MRSA, many strains exhibit resistance to additional antibiotic classes (1). The worldwide epidemic spread of MRSA is often depicted as happening in population waves caused by bursts of new, highly successful clones, which replace previously dominant ones (2). While antibiotic resistance is often invoked to explain clonal replacement in MRSA, the traits that allow an emerging lineage to succeed are rarely tested directly. Understanding the selective pressures driving strain replacement is essential to prevent the rise of new, more virulent lineages. Identifying newly emergent lineages as they first become prevalent in a region remains key to anticipating their clinical and epidemiological impacts.

Although the molecular epidemiology of MRSA is heterogeneous across South America, whole genome sequencing (WGS) has enabled identification of several strain replacement events. This is typified by the replacement of the Chilean/Cordobés (ChC) ST5-SCC*mec-*I clone by MRSA USA300 Latin American variant strains (USA300-LV, first reported in 2006) in Northern South America (3–7). The dominant clade of USA300-LV strains, referred to as the South American epidemic (USA300-SAE) clade, is a sister clade to North America’s community-associated (CA) USA300-NAE, both of which belong to sequence type 8 (ST8) (8). USA300-SAE appears to have spread quickly in northern South America in the early 2000s, replacing ChC in the northern countries of Colombia, Venezuela, and Ecuador (3,6–11). Like USA300-NAE, USA300-SAE possesses a pathogenicity island (SaPI5) carrying the enterotoxins *sek* and *seq* and genes for the Panton-Valentine leukocidin (PVL) genes, but it harbors a copper and mercury resistance (COMER) element in the place of USA300-NAE’s arginine catabolic mobile element (ACME)(8). Despite its proliferation in Venezuela, Ecuador, and Colombia, USA300-SAE has only made up a minor proportion of MRSA found in more southern countries such as Peru, Brazil, Argentina, and Chile(12). Instead, Chilean MRSA continued to be dominated by the hospital-associated (HA) ST5 ChC clone (13,14). However, Aguayo-Reyes et al. (2025) recently observed a high predominance of USA300-SAE among a sample of 54 community-associated MRSA isolates ranging from 2007 to 2024 in Chile. In a recent prospective study of combined HA- and CA-MRSA in Chile, we observed a steep decline in ST5, leading us to hypothesize that other sequence types, such as ST105 or ST72, may be in the process of replacing the dominant ChC clone in the region (13). Identifying the reasons for MRSA strain replacement is nontrivial: the spread of new epidemic strains is often attributed to antibiotic use and misuse, but other fitness advantages such as resistance to environmental selective pressures (e.g. heavy metals) or host immune response may also be at play (5,11, 22–24). Resolving whether such fitness advantages could contribute to strain replacement requires genomic resolution capable of distinguishing closely related sub-lineages.

WGS is crucial for tracking novel strains of MRSA. Molecular techniques such as *spa*-typing or multi locus sequence typing (MLST) have been useful to track major epidemiological changes but lack the granularity to identify individual clones. By contrast, whole genome sequences provide very high resolution and enable more confident reconstructions of their evolutionary pathways and relationships(15,19,20). WGS-based surveillance can track MRSA spread in large geographic regions and even uncover transmission events between people or locations (3,21,22). However, whole genome sequencing remains expensive and impractical at scale and in resource limited settings. We have recently developed several tools that use WGS phylogenetic inference to identify diagnostic (apomorphic) loci for clades of interest, leveraging the granularity of WGS to design inexpensive, scalable molecular tests (9,23–25).

Here we sought to determine how USA300-SAE emerged and expanded in Chile and whether traits beyond antimicrobial resistance could plausibly contribute to its success. We collected, sequenced, and analyzed 274 invasive MRSA isolates of both hospital and community origin observed in 2022 in a multicenter network of 11 hospitals distributed across Chile as part of an ongoing multicenter surveillance effort. We compared our 2022 cohort to previously collected MRSA from invasive infections spanning 1999 to 2018 to characterize changes in antibiotic resistance prevalence and sequence type composition of MRSA over time. We report a marked increase of ST8 over time, with a particular increase in the previously rare USA300-SAE lineage, amplifying recent observations (26). We present and validate a rapid, direct-from-colony restriction digest-based diagnostic test for USA300-SAE which we used to estimate its 2023 prevalence. We reconstructed the temporal and geographical origins of USA300-SAE in Chile and combined *in vitro* and *in silico* methods to evaluate the potential role of USA300-SAE’s mobile copper genes (*copX-mco-copL)* in enabling its expansion. We show that USA300-SAE strains are resistant to copper toxicity at levels found in tap water, and we present the first data showing that the copper resistance locus in USA300-SAE is required for this phenotype.

## RESULTS

### Temporal analyses of MRSA diversity in Chile reveals a shift in antibiotic resistance profiles and a dramatic rise in ST8 frequency

We sequenced and analyzed 274 genomes from isolates collected in 2022 from 184 invasive infections treated at 11 hospitals in Chile (Figure 1). To gauge relative abundances of strains and antimicrobial resistance profiles we deduplicated genomes by patient, taking only the first isolate sequenced from each patient. Historical comparisons of the 2022 sample to MRSA recovered from invasive infections in Santiago from 1999 to 2018 demonstrated a clear shift in recent antibiotic susceptibility profiles (13). Most notably, gentamicin, clindamycin, ciprofloxacin, and erythromycin resistance declined in prevalence among invasive MRSA infections, with gentamicin’s decline being the most dramatic (95.2% in 1999-2002 to 24.5% in 2022, Figure 1A).

**Figure 1.**
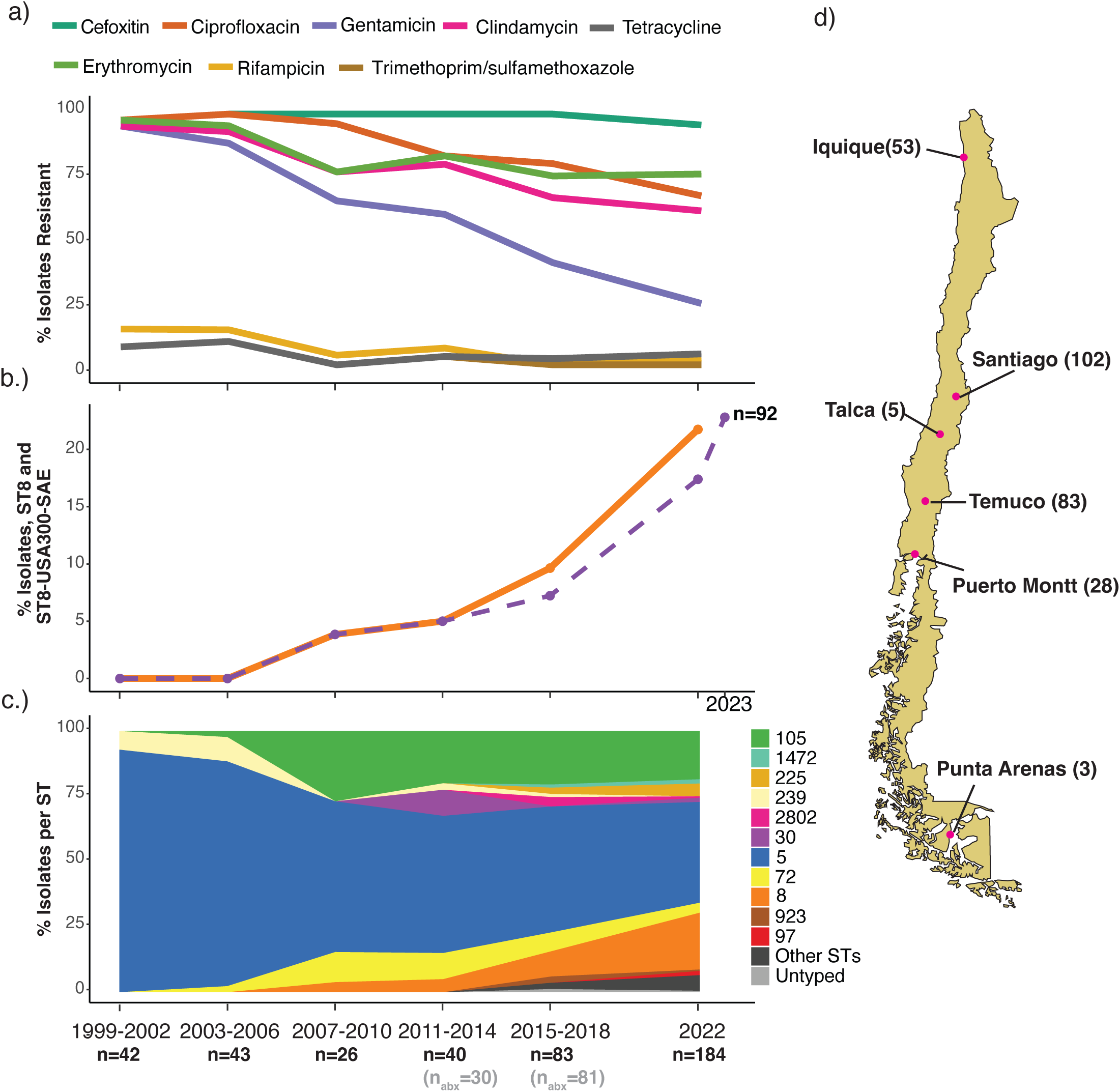
Temporal analyses of MRSA diversity in Chile show changing antibiotic resistance profiles and a dramatic rise in ST8. a-c) % isolates by year, deduplicated to one per patient a) resistant to 8 antibiotics based on disk diffusion assays b) belonging to ST8 and USA300-SAE, calculated from whole genome sequences from 1999-2022 and from molecular diagnostic tests in 2023 and c) belonging to each ST, where “Other STs” refer to STs with only one observation. d) Total number of isolates sequenced from each city in 2022

We next compared the multi-locus sequence type (ST) composition of the 2022 sample to that of previous years (Figure 1B-C). Of the deduplicated isolates from 2022, 38.6% were ST5, 21.7% were ST8, and 18.5% were ST105. We observed a surprising increase in ST8’s prevalence over time, with a particular increase between 2015 and 2022 (9.6% and 21.7%, respectively). This rise in ST8 was mirrored by the previously reported decline in ST5, which decreased from over 90% during the 1999-2006 period, to less than 40% in 2022 (13). ST105, which increased in prevalence between the 2003-2006 and the 2007-2010 periods, has stabilized at around a 20% prevalence.

These changes track with the resistance profiles we observed for the isolates belonging to each ST of MRSA tested; while ST5 was most commonly resistant to ciprofloxacin, gentamicin, clindamycin, and erythromycin, ST105 was resistant to all the same drugs as ST5 except gentamicin, and ST8 was typically not resistant to any of the tested drugs except cefoxitin, as expected (Figure S1). The decline in resistance to gentamicin, ciprofloxacin, clindamycin, and erythromycin, therefore, coincides with the decline in ST5 over time. As ST5 and its associated antimicrobial resistance profile declines in this region of South America, it is crucial to identify which strains may be taking its place. Though multiple different sequence types have changed in prevalence in the past 20 years, ST8 shows the steadiest, most consistent increase.

### Chilean ST8 MRSA is genomically diverse but is dominated by the USA300-SAE epidemic clade

We next asked whether the marked increase in ST8 in Chile from 1999 to 2022 (from 0% to 21.7% relative abundance) was due to the emergence of a specific, geographically localized lineage within the sequence type. To evaluate the diversity of Chilean ST8 in the context of the sequence type’s evolutionary history and distribution, we built a maximum likelihood phylogeny combining our Chilean ST8 genomes from 1999 to 2022 with ST8 genomes included in Bianco et al. (2023) and the more recently reported ST8 genomes from Di Gregorio et al. (2023) from Argentina, Bolivia, Brazil, Paraguay, and Uruguay (Figure 2) (22).

**Figure 2.**
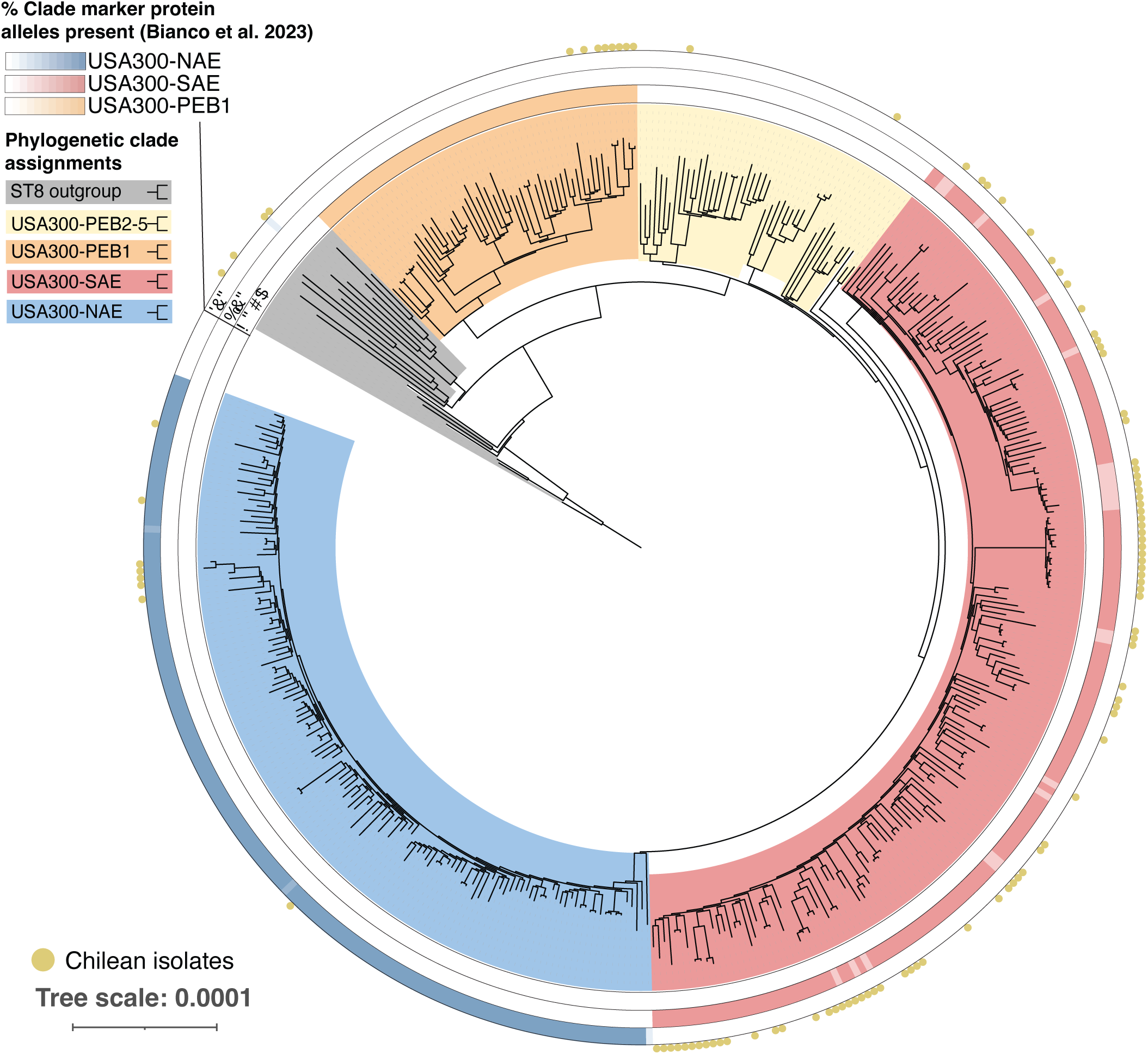
**Maximum likelihood phylogeny of 433 ST8 genomes.**Yellow circles indicate isolates collected in Chile from 1999-2022. Fill color indicates phylogenetically assigned ST8 subclade. Heatmaps indicate the % of Bianco et al. (2023) protein alleles for each subclade present by genome.

Chilean isolates showed substantial phylogenetic diversity, forming 49 monophyletic lineages, including singletons, which spanned the entire tree (Figure 2). However, 36 of these monophylies (73.4%) belonged to the USA300-SAE clade, indicating that this is the dominant clade of ST8 MRSA in Chile. Consistent with previous reports, all USA300-SAE isolates circulating in Chile harbored SCC*mec*-IVc (8). Chilean isolates were dispersed across the USA300-SAE subtree, suggesting multiple independent introductions of SAE to Chile rather than the fixation and proliferation of a single clone.

### Phylogenetically-based molecular diagnostic tests enable rapid identification of USA300-SAE and demonstrate its ongoing proliferation

In addition to classifying the 2022 Chilean ST8 isolates based on their phylogenetic placements, we also classified them using previously derived diagnostic alleles for ST8 lineages (Figure 2, Table 1). Bianco et al. (2023) reported protein alleles with high specificity and sensitivity to genomes in the USA300 (NAE and SAE), USA300-SAE, USA300-NAE, and PEB1 clades. The alleles showed very high sensitivity and specificity in a test set that combined the Chilean ST8 genomes and the Di Gregorio et al. (2023) genomes (Table 1). All diagnostic alleles for USA300, USA300-SAE, the USA300-SAE/NAE combined clade, and PEB1 showed 100% specificity for their phylogenetic placements. Four out of five diagnostic alleles for USA300-NAE were 100% specific, with the fifth being 99.4% specific. The diagnostic alleles also showed relatively high sensitivity, with all but one of them exhibiting sensitivity values > 96%. A marker allele in the *vwb* gene for USA300-SAE displayed the lowest sensitivity at 82.5%.

**Table 1.**
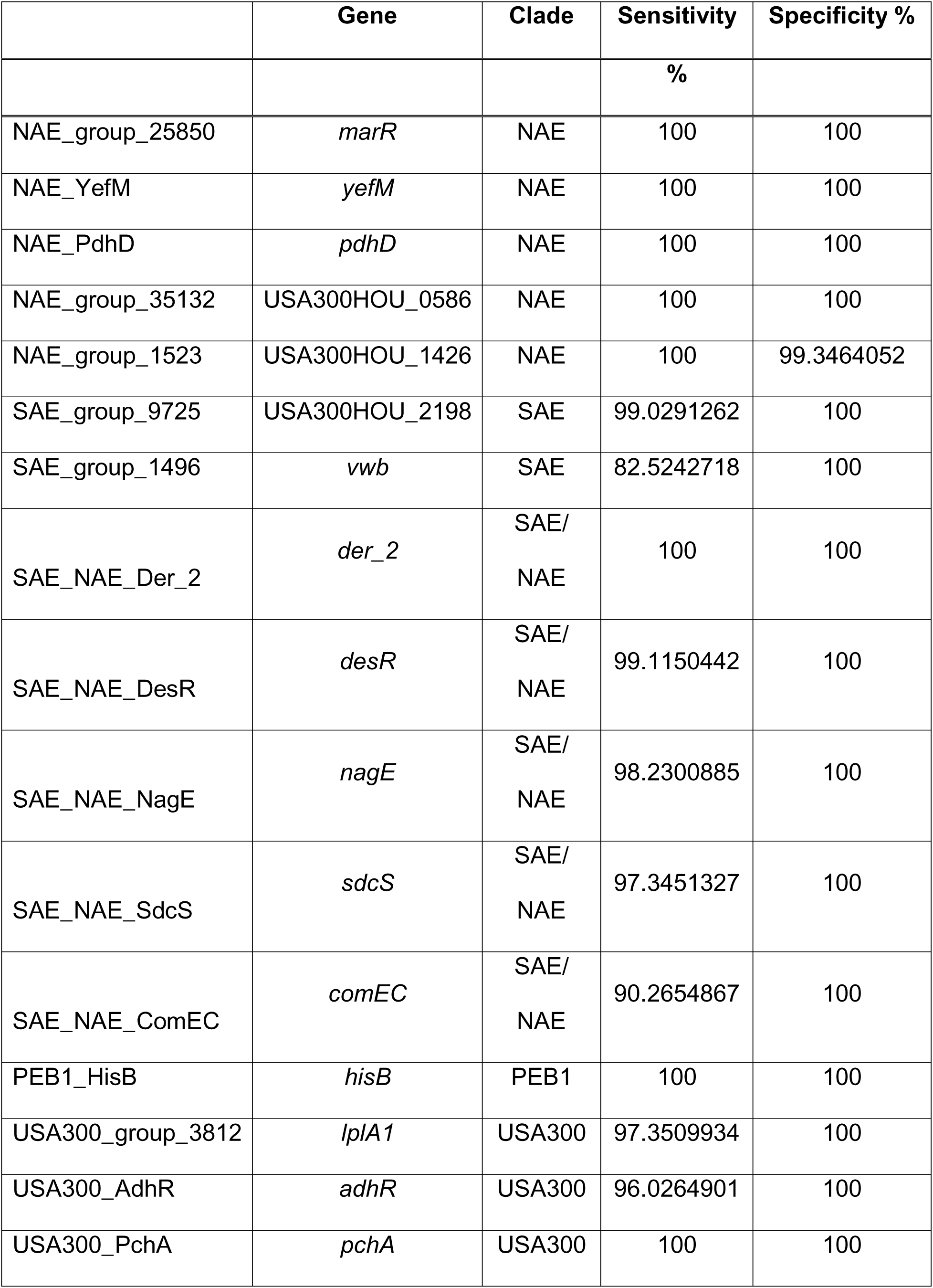
Sensitivities and specificities for Bianco et al. (2023) alleles in Chilean isolates (1999-2022) and StaphNET isolates belonging to ST8 (n=163)

With the goal of developing rapid, inexpensive molecular diagnostic methods to track the emergence of USA300-SAE in real time, we used two whole-genome sequencing-independent methods to calculate the prevalence of USA300-SAE among invasive MRSA isolates collected in 2023. First, we used Classification Using Restriction Enzyme Diagnostics (CURED), a new method which generalizes Esteves et al. (2023)’s restriction endonuclease test by identifying high-specificity and high-sensitivity k-mers for phylogenetically determined groups of interest and identifying restriction enzyme sites within diagnostic k-mers (23,25). Using USA300-SAE genomes from Figure 2 as ‘cases’ and a combination of non-USA300-SAE ST8 genomes from Figure 2 and non-ST8 genomes from Chile in 2022 as ‘controls,’ CURED identified 32 k-mers with 100% sensitivity and 99% specificity to USA300-SAE. The 99% specificity was due to 3 control genomes which were in fact single locus variants of ST8 and grouped phylogenetically in the USA300-SAE clade. Thus, the actual sensitivity and specificity of the k-mer to the USA300-SAE clade were 100% and 100%. Of these 32 k-mers, 9 had restriction enzyme recognition sites (Table 2). We designed a BsmAI restriction enzyme digest of the region including one diagnostic k-mer which yielded 396-length and 140-length fragments if the amplified region included the BsmAI site, and a single 536-length fragment if it did not (Figure S3).

**Table 2.**
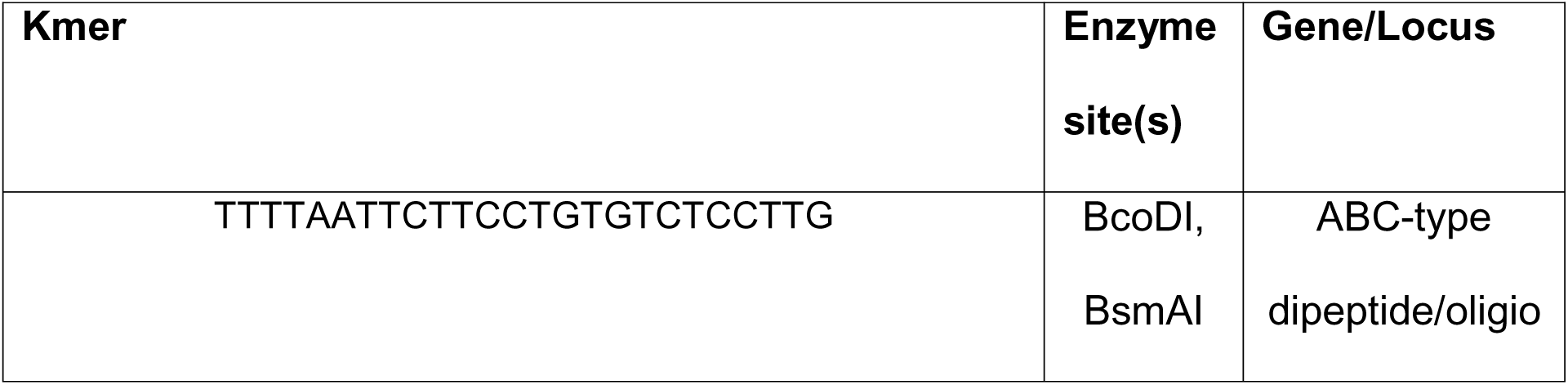
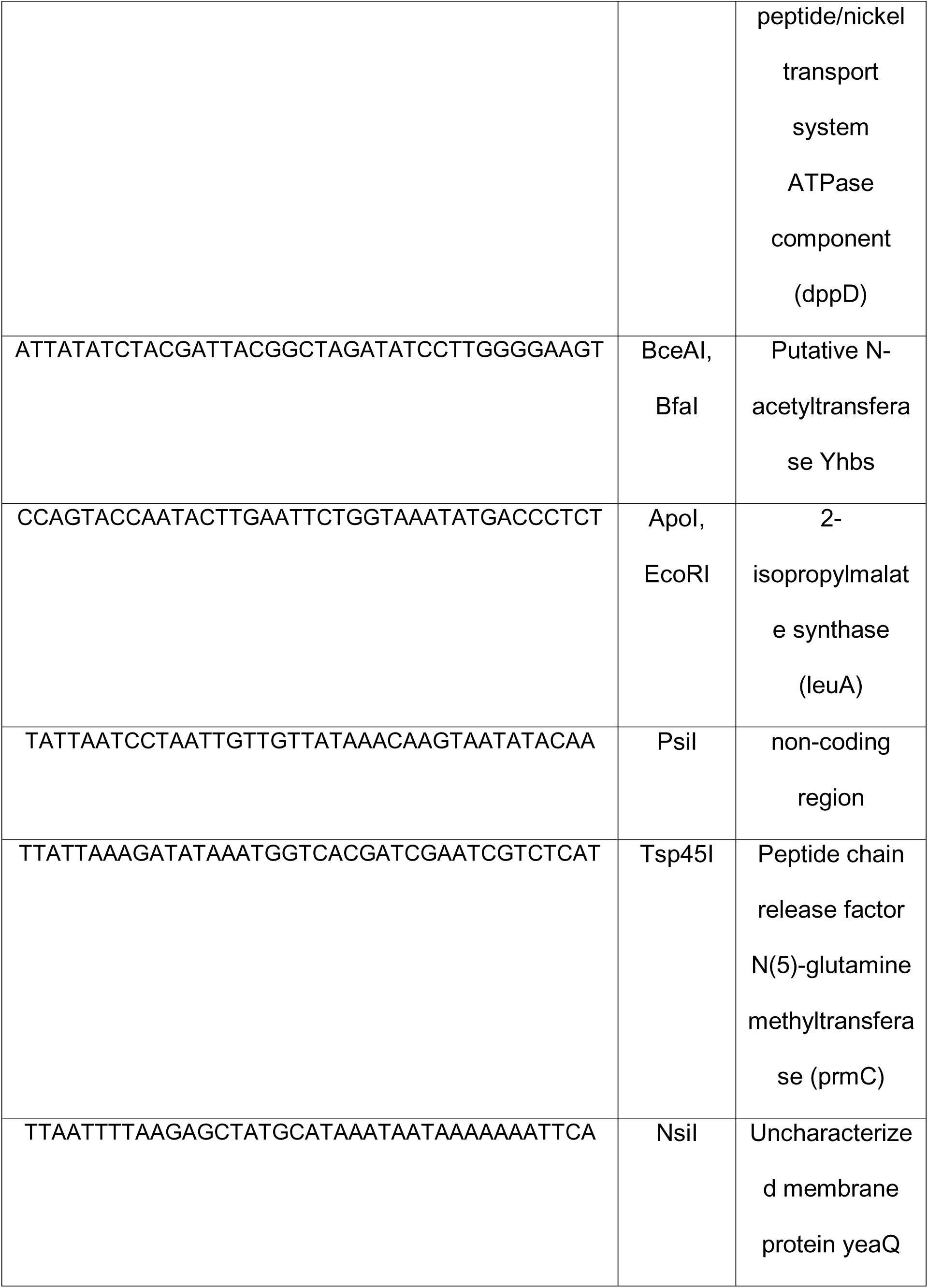
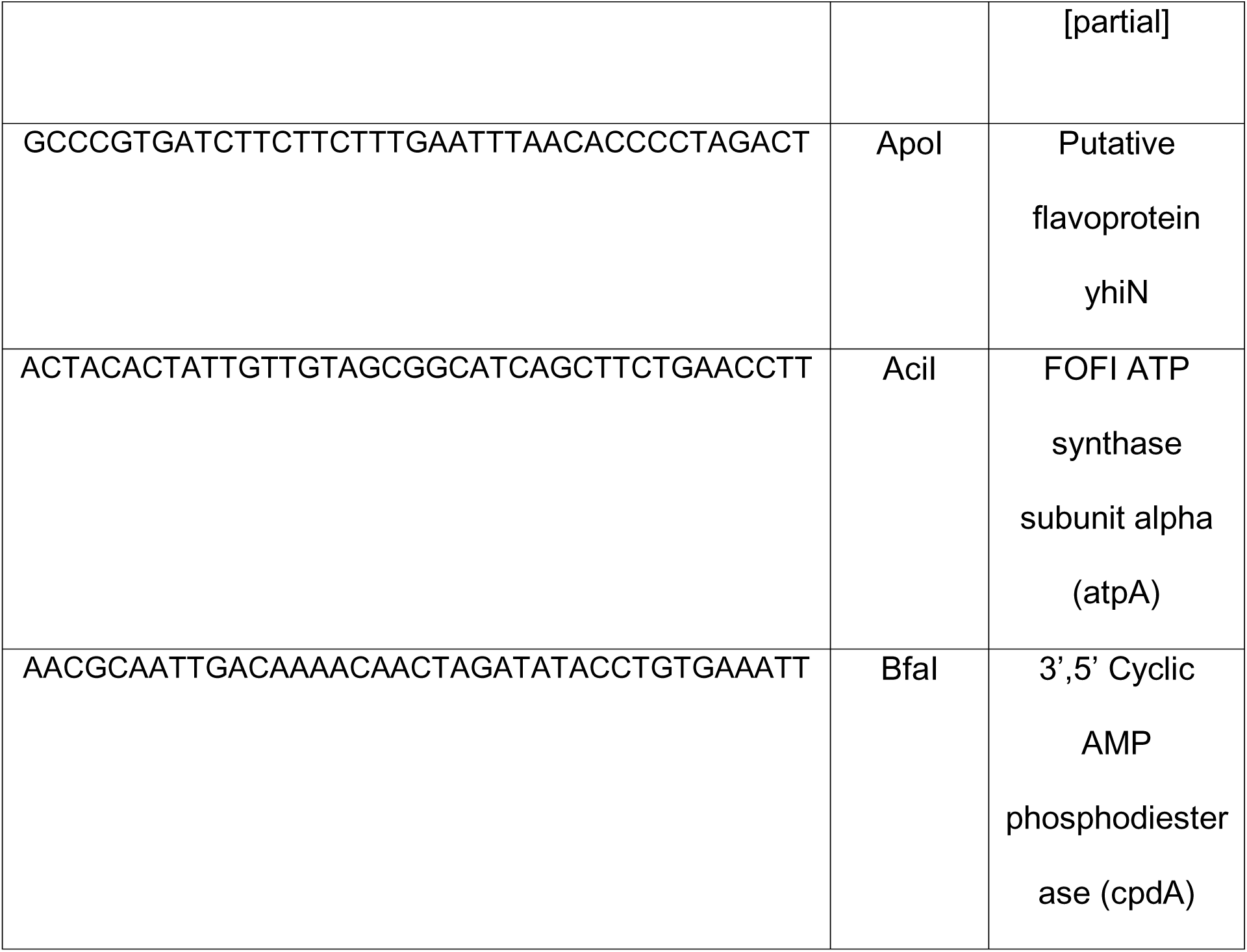
100% sensitive and 99% specific USA300-SAE unique k-mers identified by CURED containing restriction enzyme sites.

Using this CURED-based test we assayed 92 un-sequenced MRSA isolates collected from Chile in 2023. Of these isolates 21(22.8%) were classified as USA300-SAE (Figure 1B).

In a second WGS-independent approach, we cross-validated the CURED diagnostic method by performing Sanger sequencing on the strongest USA300-SAE diagnostic allele from Bianco et al. (2023), a G to A substitution resulting in the introduction of a new stop codon in the gene for hypothetical protein USA300HOU_2198 (Table 1)(9). Seventy-six isolates produced a positive band on PCR and yielded identical classifications to the CURED restriction enzyme test. The remaining sixteen isolates were predicted to be non-SAE by the CURED-based restriction and failed to consistently produce a band of the expected size with the USA300HOU_2198 primers, raising the possibility that the targeted PCR locus was absent or altered in distantly related strains.

Whole genome sequencing of these 16 isolates showed that one ST97, seven ST72 and two ST1649 genomes had a 2kb deletion including the USA300HOU_2198 gene and the two genes flanking it. The remaining four ST398 genomes, one ST101 genome, and one ST25 genome contained most of the diagnostic PCR product but lacked part or all the reverse primer target sequence. In total, results for the restriction enzyme-based approach were 100% concordant with their results from the HOU_2198 allele sequencing-based approach when the HOU_2198 gene was present in the genome.

We validated our approach using whole genome sequencing of a set of 10 predicted SAE and 10 predicted non-SAE isolates. Phylogenetic inference of these genomes showed that our molecular diagnostic methods were 100% accurate for this subsample (Figure S4B). These highly accurate molecular diagnostic methods, which enable rapid strain typing of US300-SAE without high-throughput sequencing, indicate that the prevalence of USA300-SAE rose from 17.4% in 2022 to to 22.8% in 2023. This represents a continued rise in USA300-SAE prevalence beyond our 2022 sequencing cohort, consistent with the upward trend observed between 2015 and 2022 (Figure 1).

### USA300-SAE was introduced to Chile multiple times from other countries

To infer the temporal and geographic origins of USA300-SAE in Chile, we performed a Bayesian molecular clock analysis and geographic ancestral state reconstruction on a tree of 155 USA300-SAE genomes with known countries and dates of collection, which we rooted at *S. aureus* genome HUV01, an early branching USA300 isolate found outside of the USA300-SAE and NAE clades (Figure 3) (11). We identified 6 different locally circulating Chilean USA300-SAE sub-lineages, defined as clades with a predicted Chilean ancestor (marginal posterior probability > 50%) containing Chilean isolates from multiple different patients. The predicted divergence dates of the six locally circulating sub-lineages ranged from 2001 (95% HPD=[1999.9, 2002.6]) to 2019 (95% HPD=[2018.6,2020.5]).

**Figure 3.**
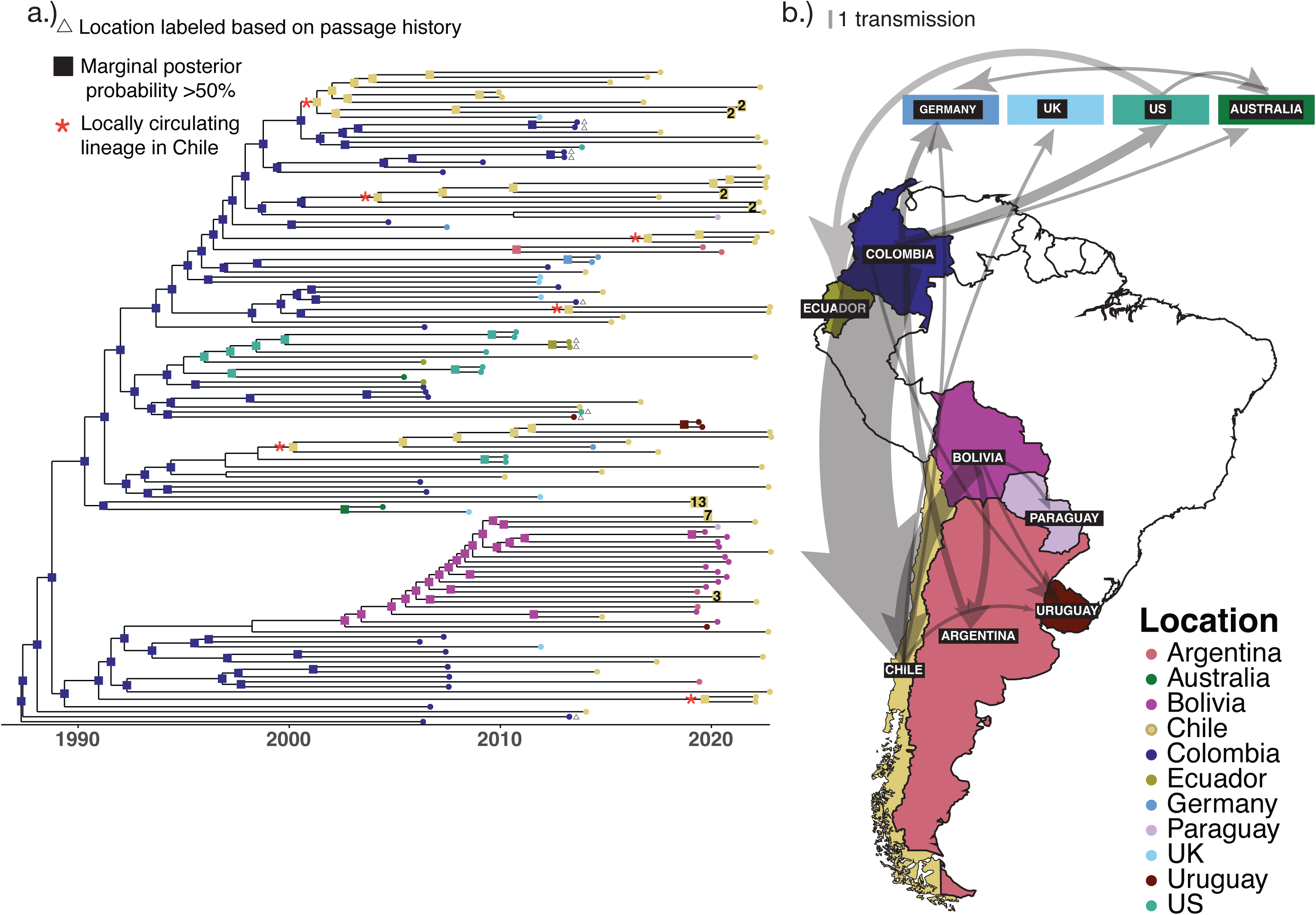
Temporal and geographic dynamics of USA300-SAE spread. a) Phylogeny of 155 USA300-SAE genomes with molecular clock estimates (timescale) and ancestral reconstruction of location. Nodes with >50% marginal posterior probabilities for one country are labeled. Boxed numbers indicate the number of genomes collapsed into a monophyletic clade within patient. b) Inferred transmission map based on changes in location between adjacent nodes in (a). Arrow line thickness is proportional to the number of inferred transmissions.

Although most Chilean clades and singletons were predicted to originate in Colombia, one clade that grouped 6 Chilean isolates with the Bolivian USA300-SAE genomes reported in Di Gregorio et al. (22) was predicted to originate in Bolivia in 2002 (95% HPD=[2000.9, 2004.2]). Rather than forming a single monophyletic clade within the Bolivian clade, Chilean genomes were interspersed with genomes from Bolivia, Paraguay, Uruguay, and Argentina. Chilean isolates from this clade were largely localized to the North of the country: 4 out of 6 of its Chilean isolates were isolated in the northern city of Iquique, which is less than 100 miles from the Bolivian border. The ancestral reconstruction also suggests transmission of USA300-SAE from Chile to other countries. For example, a small clade of two Uruguayan isolates collected in 2019 were predicted to have a recent Chilean ancestor. Another member of the same locally circulating Chilean clade from 2000 (95% HPD = [1998.2, 2002.0]) was isolated from Germany in 2014 (27). Together, these results indicate that USA300-SAE has been introduced into Chile multiple times over the past two decades, from different geographic origins.

### Recurrent, independent loss events in COMER consistently preserve the copper resistance cluster

The copper and mercury resistance element (COMER) is a defining feature of the USA300-SAE, yet its contribution to the lineage’s epidemiological success remains untested. Given USA300-SAE’s limited antibiotic resistance profile (Figure S1), we investigated whether COMER’s copper resistance genes could provide an alternative fitness advantage underlying its expansion in Chile. COMER, as described in Planet et al (8), contains three distinct gene clusters: a cluster of phage abortive infection related genes (*abi* genes) associated with SCC*mec-*IVc, a cluster of mercury resistance genes, and a cluster of copper resistance genes designated *copX* (also referred to as *copB*), *mco*, and *copL*(8,28,29). We identified 14 different loss events affecting portions of COMER across the tree of USA300-SAE genomes included in the molecular clock and ancestral reconstruction analysis (Figure 4A). The copper resistance gene cluster was more highly conserved than the rest of the COMER element and was affected by only 4/14 loss events compared to 10/14 affecting the mercury resistance cluster and 9/14 affecting the *abi* cluster. Notably, we observed convergent losses of the mercury resistance and *abi* gene clusters that left the copper gene cluster intact, including a subclade of the Bolivian clade identified in the USA300-SAE phylogeny. Moreover, we observed the same region lost in phylogenetically independent isolates from two additional patients from the 2022 Chilean cohort (Figure S3A).

**Figure 4.**
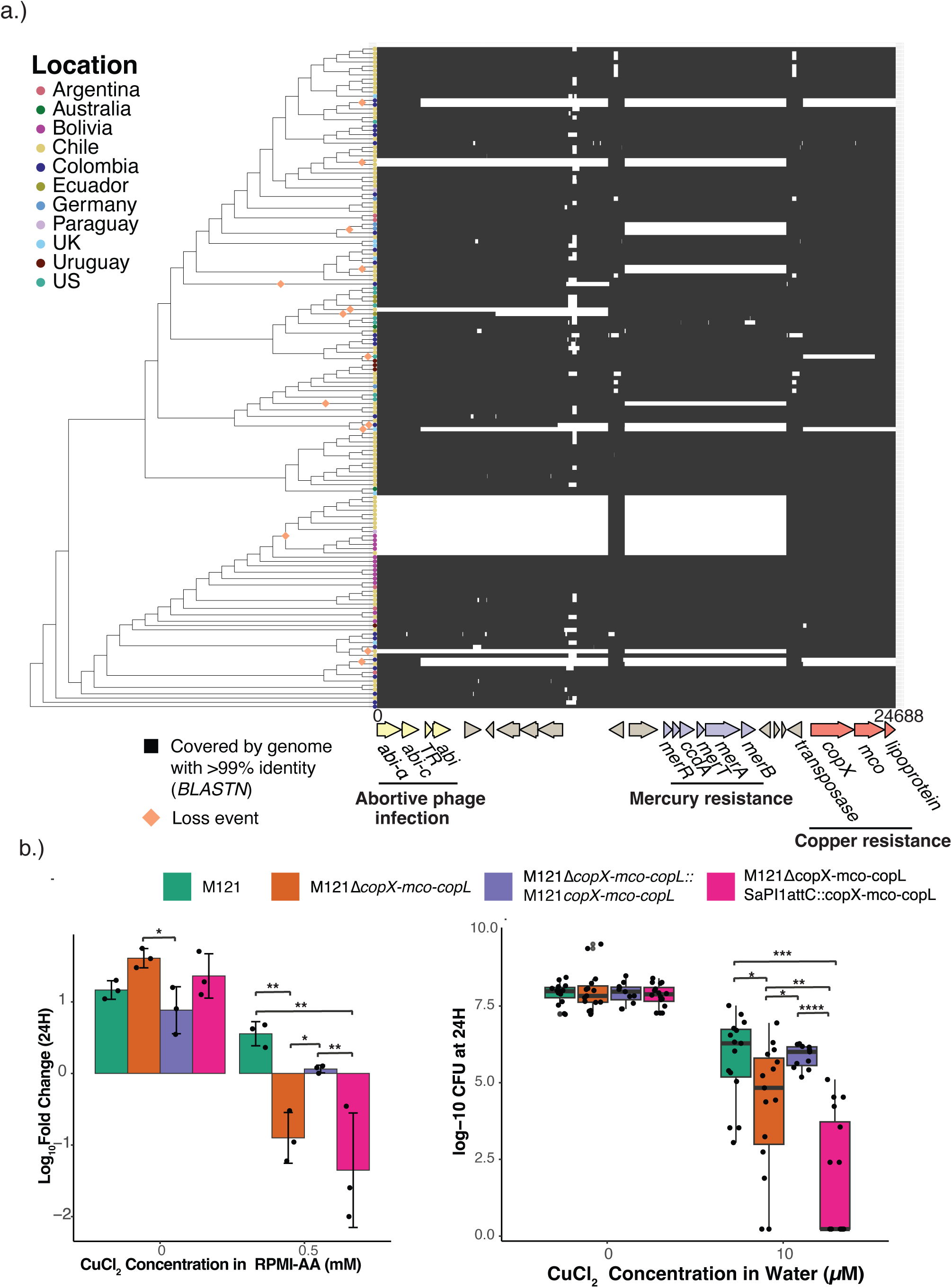
Conservation of COMER-conferred copper resistance in USA300-SAE. a) Coverage of COMER element among USA300-SAE. Loss events are highlighted for losses >1 KB. b) Protection against copper in RPMI-AA media(left) and water (right) by COMER gene cluster *copX-mco-copL.* Significance is reported for wilcoxon(right) and t-tests (left) corrected for multiple comparisons via the Benjamini-Hochberg process where * < 0.05, ** < 0.01, *** < 0.001, ****<0.0001

These observed gene losses also affected the SCC*mec* element, which is located immediately adjacent to COMER and harbors the *mecA* methicillin resistance gene. Based on alignments of genomes’ sequencing reads to a SCC*mec* IVc/COMER reference sequence, we observed that all genomes missing the most proximal part of COMER to SCC*mec* were also missing part of SCC*mec* (Figure S4A). Two of the three loss events resulted in the loss of COMER with preserved copper genes started at the same position, ranging from immediately upstream from SCC*mec*’s *ccrA/ccrB* to immediately upstream of the transposase adjacent to *copX*. The third loss event corresponded to a range starting in the middle of *mecR1* and again ending upstream of the copper cluster-adjacent transposase. A fourth SCC*mec*/COMER loss event, affecting the *abi* cluster (associated with the SCC*mec*IVc element) but not the adjacent mercury resistance genes, ranges from immediately downstream of the ISSau3 transposase of SCC*mec* to immediately upstream of the transposase adjacent to the mercury cluster.

### COMER’s *copX-mco-copL* locus confers resistance to copper in water and media

Given the broad retention of the copper resistance cluster despite recurrent loss elsewhere in COMER, we asked whether this cluster provides increased copper tolerance to USA300-SAE. The genes *copX* and *copL* in USA300-NAE’s ACME element have previously been shown to increase survival and growth in defined media with copper compared to deletion mutants for these genes(29,30). However, the same has not been shown for COMER’s *copX-mco-copL* locus. To evaluate whether this gene cluster reduces copper toxicity in USA300-SAE, we deleted the copper locus in the USA300-SAE strain M121 (Δ*copX-mco-copL*). When grown in the defined media RPMI-AA with a range of copper concentrations we saw significant protection from copper toxicity from 0.5mM to 4mM with the strongest protection at 0.5mM (Figure 4B, Figure S4A) This phenotype was complemented by reintroduction of the entire *copX-mco-copL* locus in cis as a single copy(31), demonstrating that this locus is required for the copper resistance phenotype.

We hypothesized that the copper resistance locus may also protect against copper exposure in environmental contexts. To test this hypothesis, we developed an assay that exposed the mutant and wild type to deionized water for 24 hours with varying concentrations of copper (II) chloride (CuCl_2_). We observed the most consistent difference in survival at 10µM CuCl_2_, a concentration of copper near those seen in tap water (32). Protection against 10µM copper in water was also complemented by re-introducing the *copX-mco-copL* locus in cis to the M121Δ*copX-mco-copL* mutant.

Altogether, in addition to being highly conserved over the history of USA300-SAE, the copper resistance genes of COMER protect bacteria against copper toxicity that impedes growth in culture and causes death in an aqueous, nutrient-limited context, such as might be seen on environmental surfaces.

## DISCUSSION

The worldwide dissemination of MRSA is often characterized by different lineages dominating in different geographic areas. During the past two decades, the MRSA epidemiology in Chile has been marked by a clear dominance of the healthcare-associated ST5 ChC clone of MRSA (7,10,13,14). During this time, the South American Epidemic clade USA300-SAE, which is closely related to the North American epidemic clade USA300-NAE, became widespread in Colombia, Venezuela and Ecuador, replacing the dominant ST5 ChC clone in these countries.(7,9,11,33). As USA300-SAE took over in Northern South American countries, it remained rare in Southern South American countries including Chile, Peru, Brazil and Argentina (3,13).

We now present evidence that USA300-SAE is taking hold as a dominant MRSA lineage in Chile after numerous different introductions. Our data show that a consistent increase in ST8 prevalence accompanied a decline in ST5 in Chile between 2000 and 2022, and that this emerging ST8 predominantly belongs to the USA300-SAE clade. Using a molecular diagnostic method, we showed that USA300-SAE was of even higher prevalence in 2023 compared to 2022, indicating that it is still likely expanding in the region. These findings are consistent with recent reports of USA300-SAE infections at a hospital in Concepción in South-Central Chile, and a subsequent analysis of 54 CA-MRSA isolates showing its dominance among community-associated cases of MRSA in Chile (26,34).

We previously reported increased ST diversity among invasive MRSA in Santiago between 2000 to 2016, and hypothesized that ST72 and ST105 could be replacing the ST5 ChC clone in Chile (13). While ST105 remained prevalent at 18.5% relative abundance in our new data from 2022, ST8 surpassed it at 21.7%. Combined with ST8’s steady increase over recent years, these new data suggest that ST8, rather than ST105 or ST72, is in the process of replacing ST5 as the dominant clone in this part of South America.

The decline in ST5 was accompanied by a decrease in antibiotic resistance among invasive MRSA in Chile, largely reflecting the loss of the highly resistant ST5 ChC clone, which was frequently resistant to gentamicin, ciprofloxacin, clindamycin, and erythromycin (Figure S1). It is possible that the ST5 decline and concomitant rise in more susceptible MRSA lineages such as ST8 is due to its broad antibiotic resistance: though antimicrobial resistance genes can enable certain strains to persist in a population on which their target antibiotics are used, they can also carry a fitness cost when not strictly necessary for survival (35–37). However, our correlative data cannot infer such a causal relationship, and further investigation into the relative decrease of ST5 prevalence over time is required. Still, the question remains as to why ST8, which in our collection is resistant to only beta-lactam antibiotics, is rapidly increasing in relative abundance among invasive MRSA in Chile.

The decline in ST5 ChC prevalence in Chile predates the expansion of ST8, beginning as early as 2007 (13). Moreover, prior to the 2015-2018 period, ST105 and ST72 were far more prevalent than ST8. It seems most likely, therefore, that ST8 is replacing an ST5 population that is declining due to other factors, rather than overtaking and displacing it. Strain replacement in MRSA is often attributed to antibiotic use, which can select for strains with chromosomally or plasmid-encoded antibiotic resistance (2). The initial emergence of MRSA, for example, was associated with early beta-lactam use selecting for the *mecA* gene (38). However, outbreaks associated with increased incidence of disease are often attributed to other survival phenotypes, including those related to interactions with the host environment. The spread of the USA300-NAE clade of MRSA was immediately preceded by its acquisition of the ACME genetic element, which is thought to enhance survival on skin, while the SCCmec element conferring methicillin resistance was acquired much earlier (9,16,17). In Europe, ST22-IV most likely replaced ST228-I due to its resistance to environmental and host stressors, rather than antibiotic selection (15,18). It is therefore plausible that environmental or host factors, rather than antibiotic pressure, underlies USA300-SAE’s recent success in Chile.

One possible explanation for the expansion of USA300-SAE despite its relatively limited antimicrobial resistance arsenal is its metal resistance profile. Chile is the largest producer of copper in the world and has especially dense copper deposits and mining operations in the northern half of the country (39). Both arsenic and copper are found in relatively high concentrations in groundwater in Chile due to a combination of natural geologic and industrial factors (39,40). Since arsenic and copper are toxic to *S. aureus* via their catalysis of redox reactions, it is plausible that a MRSA strain’s relative fitness in Chile could depend on its ability to resist metal-induced stress (41–43).

Consistent with this potential explanation, we observed broad conservation of mobile element-encoded copper resistance genes in USA300-SAE. Located immediately downstream from the SCC*mec* element, the COMER element was acquired by the common ancestor of the USA300-SAE clade independently of USA300-NAE’s acquisition of ACME and is unique to USA300-SAE among ST8 strains (8,9,44). Though some members of USA300-SAE lack some portions of COMER, complete loss of the element is rare (9,33). We found that the copper cluster *copX-mco-copL* is frequently maintained even when other parts of COMER are lost, suggesting that they may confer a selective advantage to members of this successful clade. We demonstrated *in vitro* that *copX-mco-copL* confers protection to USA300-SAE against copper in both defined media and in water. While the high concentrations of copper needed to see differential toxicity in media may or may not be physiologically relevant, the levels in our water killing assay are in line with concentrations seen in tap water(45), to which *S. aureus* strains colonizing the skin or surfaces in the environment are likely exposed. Together, these findings suggest that environmental copper exposure could have contributed to the success of USA300-SAE, though establishing this link will require evidence beyond the correlative and *in vitro* data presented here. It is also possible that the copper gene cluster in COMER strengthens USA300-SAE’s invasive phenotype: USA300-NAE’s *copX* and *copL* have previously been shown to promote infection and enable survival in macrophages (30,46).

Given the relatively recent rise of USA300-SAE in Chile, we wondered where and when the strain was introduced: did a single sub-lineage of USA300-SAE arrive and proliferate to create a Chilean clade, or were different sub-lineages introduced on multiple occasions? Our molecular clock and ancestral reconstruction analyses suggest USA300-SAE has been independently introduced to Chile from Colombia and other countries dating as far back as 2001 and as recently as 2019. This observation stands in contrast to other instances of rapid, geographically specific expansion of certain MRSA strains. When the Rio de Janiero (RdJ) clone of ST105 was identified in Brazil, for example, it was found almost exclusively among isolates from Brazil (23,24). Our results indicate multiple different introductions of USA300-SAE to Chile.

Bolivia, which borders Chile to the Northeast, was the inferred source of at least one such introduction. Di Gregorio et al. (2023) previously reported a Bolivian cluster of USA300-SAE isolates collected in 2019 which had also been introduced to Argentina, Paraguay, and Uruguay. We found isolates from six different patients in Chile belonging to this cluster: all six were from either Iquique or Santiago, and all six were dispersed across the clade rather than forming a single Chilean subclade. This phylogenetic structure and the geographically localized sampling of this clade suggest repeated transmission of the sublineage to Northern Chile, rather than a single introduction followed by local spread. This is consistent with Di Gregorio et al.’s observation that this strain was primarily community acquired rather than hospital acquired in their collection.

In total, we observed over 50 predicted transmission events between Chile and other countries (transitions between nodes on the ancestral reconstruction from one >50% marginal probability location to another), the vast majority of which were from Colombia to Chile (Figure 3B). This transmission trend could be due to sampling bias: 81/155 genomes included in the USA300-SAE analysis were from Chile, while 25/155 were from Colombia. We observed only three instances of predicted transmission from Chile to other countries: one to the UK, one to Germany, and one to Uruguay. The presence of USA300-SAE in other continents is not unexpected: USA300-SAE and LV isolates have occasionally been reported in European and Asian countries such as Belgium, France, and Japan(47–49). Overall, frequent transmission within South America would explain the diversity of USA300-SAE we found in Chile. Isolates collected in 2022 were also not limited to a single clade or transmission event but were rather spread across the USA300-SAE phylogeny. This finding suggests that they derive from multiple different ancestral lineages rather than a single, dominant epidemic sublineage. This pattern is consistent with a model of expansion into an open niche such as that left by the decline of the previously dominant ChC clone.

Developing scalable, cost-effective methods for tracking emerging MRSA strains such as USA300-SAE is essential to responding to them effectively. We previously published phylogenetically-derived diagnostic alleles for USA300, USA300-SAE, USA300-NAE, and a pre-epidemic branching clone of USA300 (9). Here, we found that nearly all of these diagnostic alleles showed 100% specificity for their diagnostic clades among independently collected ST8 genomes. While sensitivity displayed higher variability by allele, it was still high across the board, with each clade having at least one allele with >99% sensitivity. We used sanger sequencing of one of these validated alleles, alongside a restriction digest-based approach using the recently released CURED pipeline, to rapidly estimate the percentage of 2023 invasive MRSA isolates in Chile belonging to USA300-SAE. Interestingly, we identified that the hypothetical gene (HOU_2198) used for our genotyping scheme was part of a 2kb deletion from ST72, ST97, and ST1649 genomes, highlighting a caveat of an allele presence/absence-based diagnostic test. Our results show that USA300-SAE is likely still on the rise and motivate future development of rapid, lower cost, genetic diagnostics for *S. aureus* strains of interest.

Several limitations should be considered when interpreting our findings. First, we restricted our analysis to infections clinically classified as MRSA and therefore cannot identify temporal trends in methicillin susceptible *Staphylococcus aureus* (MSSA). Second, we limited our 2022 collection and historical comparisons to MRSA cultured from invasive infections and therefore cannot make conclusions about MRSA diversity among colonizing isolates. We also did not collect detailed clinical information, precluding us from attributing sequence type composition to hospital-versus community-acquired infections with certainty. Still, our 2022 collection, to our knowledge, is the most comprehensively sampled cohort to date of MRSA in Chile. Since historical samples were limited to Santiago, we cannot make geographically specific historical comparisons in the other sampled cities. Finally, our analysis of ST and sub-ST lineage prevalence is in relative terms. Without overall disease incidence data, we cannot distinguish whether an emerging strain such as USA300-SAE is outcompeting the historically dominant strain or whether it is at the center of an outbreak which is not affecting the dominant strain’s overall incidences. Future sampling is required to confirm if USA300-SAE continues to be on the rise in 2024-2026 and should be analyzed with respect to overall disease incidence data to better distinguish these two models of strain emergence.

Whether by taking over the MRSA population as an outbreak or by filling a niche left open by the decline of previously dominant strains such as ST5, ST8 USA300-SAE is on the rise among invasive MRSA isolates in Chile. Interestingly, this clade’s rise in Chile is relatively recent, despite its multiple introductions and its presumed presence in the country for decades. Our molecular clock analysis places the earliest locally circulating Chilean sub-lineages of USA300-SAE as far back as 2001, suggesting that it was present at that time, but not rapidly expanding. Rather than taking over as the dominant clone immediately upon its introduction due to high relative fitness, it only expanded recently. This lag between introduction and expansion introduces a new question about USA300-SAE in South America: why now? One possibility is that ST5 MRSA dissipated from the region for other reasons, and USA300-SAE had a high enough relative fitness in the region to fill its niche. The COVID-19 pandemic could have contributed to this shift, as travel restrictions, social distancing, and reduced antibiotic use during 2020-2021 altered both transmission dynamics and selective pressure for broad antimicrobial resistance (50,51). However, our analyses show that USA300-SAE started to increase in prevalence prior to 2018 (Figure 1), suggesting that the COVID-19 pandemic may have exacerbated, but cannot alone explain its proliferation. Though the causes are not abundantly clear, the dominant MRSA population in Chile is transitioning from a historically hospital-associated, broadly antimicrobial resistant lineage to a more commonly drug-susceptible, community-associated lineage. These largescale population changes motivate future research into their underlying evolutionary drivers, and into the common drivers of MRSA clonal replacement.

## MATERIALS & METHODS

### Isolate collection and resistance characterization

Isolates were collected from invasive MRSA infections at 11 Chilean healthcare centers between January and December 2022 (one hospital in each of Temuco, Puerto Montt, Talca, Iquique, and Punta Arenas, as well as at five hospitals in Santiago). Isolates were sent to a reference laboratory for further characterization. where *S. aureus* species was confirmed by MALDI-TOF (Microflex LT Bruker Daltonik, Bruker, Germany). Methicillin resistance was confirmed using a cefoxitin disk diffusion assay, but isolates were included downstream if they were initially classified as MRSA in the clinic. MRSA isolates were also evaluated for susceptibility to rifampicin, erythromycin, sulfamethoxazole/trimethoprim, tetracycline, ciprofloxacin, clindamycin, and gentamicin based on additional disk diffusion assays in accordance with guidelines established by the Clinical and Laboratory Standard Institute (CLSI, 2023).

### Genome sequencing and analysis

279 isolates were streaked onto *S. aureus* ChromAGAR and single colonies were grown overnight in LB and sequenced on the Illumina NovaSeq 6000 v1.5 flow cell to produce 2 x 150bp paired reads. Raw reads were trimmed of adapters and decontaminated of host reads using *Sunbeam 4.6.0* (52). Decontaminated reads were checked for quality with *FastQC 0.12.1* and trimmed using *TrimGalore 0.6.10* (53). Processed reads were assembled into contigs using *Shovill 1.1.0* (54). Assemblies were checked for contamination using *CheckM* 1.2.2 with a contamination cutoff of 5% and a completeness cutoff of 95%, and further checked for taxonomic identity using *mash screen* against the *RefSeqSketchesDefaults.msh* database from RefSeq release 70 (55,56). Following quality control, 274 sequences were used in downstream analyses. Contigs were sequence typed using *mlst 2.23.0*, and untyped sequences were manually reviewed to include single nucleotide variants of existing STs (57,58). Sequences were deduplicated by patient for compositional analyses of relative ST abundance and antibiotic resistance prevalence. The sequence from the earliest collected isolate for each patient was selected. All ST8 genomes, regardless of patient, were included in the ST8 tree.

### Maximum likelihood phylogeny of ST8

The maximum likelihood phylogeny was constructed from a combination of genomes used in Bianco et al. (2023), genomes published in Di Gregorio et al. (2023), ST8 genomes from our 2022 collection, and Chilean ST8 genomes collected in previous years. All ST8 reference genomes were run through the same contamination and taxonomic check as the newly sequenced isolates. A pairwise multiple alignment was constructed using *Snippy 4.6.0* with *GCA_000017085.1* as a reference sequence. *RaxML 8.2.13* was used to construct the maximum likelihood phylogeny with 100 random starts, the *GTRGAMMA* model with 4 discrete rate categories, and a random seed of *19104*. *ClonalFrameML 1.12* was used to detect regions of recombination based on the resulting phylogeny and mask them from the alignment using the *- output_filtered* parameter (59,60). The filtered alignment was then used to rebuild the maximum likelihood phylogeny with the same parameters as before, and support was measured with 100 bootstrap pseudoreplicates. Outlier detection on an early iteration of the tree using *TreeShrink 1.3.9* identified ERR9884500 for removal from the analysis.

### ST8 subtyping and allele sensitivity/specificity analysis

All included ST8 genomes were annotated using *Bakta 1.8.2* and their resulting predicted amino acid CDS were checked for exact matches to the diagnostic protein alleles reported in Bianco et al. (2023) (61). Clade assignments were confirmed based on genomes’ placement on the maximum likelihood phylogeny. Sensitivity and specificity for each allele was calculated for a test set of all Chilean ST8 genomes and all genomes from Di Gregorio et al. (2023) genomes. SCCMec types were characterized using *Staphopia’s sccmec* method, and untyped genomes were further reviewed using *SCCmecFinder* and *blastn* searches for individual *SCCmec* genes.

### *In vitro* molecular typing: CURED

To track the emergence of USA300-SAE in real time, we used the Classification Using Restriction Enzyme Diagnostics (CURED) pipeline. (25)The case group consisted of 174 USA300-SAE genomes from Figure 1C, while the control group included 465 non-USA300-SAE ST8 genomes from Figure 1C as well as non-ST8 genomes from Chilean isolates collected in 2022. K-mers specific to the case group were identified via the *CURED_Main.py (v1.0)* script and filtered to those present in the fewest control genomes, yielding a total of 32 k-mers with >99% specificity to the case group. The three genomes which broke the 100% specificity threshold were subsequently shown to be single nucleotide variants of ST8 which grouped phylogenetically with USA300-SAE, making the true specificity of the 32 k-mers 100%. The *CURED_FindREs.py (v1.0)* script identified unique restriction enzyme sites in 9 of the 32 case-specific k-mers, and custom scripts available in https://github.com/amyecampbell/ChileUSA300. The 9 k-mers were subsequently queried against 79,921 publicly available *S. aureus* genomes (62).

### *In vitro* molecular typing: rapid restriction digests for diagnostic alleles

We selected k-mer TTTTAATTCTTCCTGTGTCTCCTTG which includes the unique restriction site for BsmAI, GTCTCC. We designed primers to amplify a 536-length sequence including the k-mer using *S. aureus* M121 as a reference for USA300-SAE (**Forward**: 5’-CCTTTGATATTACAGCAGGG-3’, **Reverse**: 5’-CTGCATCGTTACGTCTAATG-3’), which we amplified direct from colony using GoTaq Green (*Promega* Cat no. M7123) in a 20µL reaction. The PCR product was then directly digested with 5µL 10X rCutSmart buffer (*NEB* B6004S) combined with 1µL BsmAI (*NEB R0529S)* for 90 minutes at 37°C. The digested product was run on a 1.5% agarose gel at 100V. This protocol was carried out in 96-well plates including 46 unknown 2023 isolates in duplicate as well as previously sequenced positive and negative controls (ST8-USA300-SAE M121, internal ST5 PP.3350). Isolates that yielded two bands of sizes ∼396 and 140 in at least one replicate were classified as USA300-SAE.

### *In vitro* molecular typing: sanger sequencing of diagnostic HOU_2198 locus

We designed primers to amplify a 497-length insert (**Forward:** 5’ CAATAGCATACCTTTGATGTG-3’, **Reverse:** 5’-GTGTAGACGATTCAGGTAGC-3’) including the G195A SNP responsible for the HOU_2198 diagnostic allele for USA300-SAE published in (9). We did sanger sequencing using the forward primer, and classified sequences as USA300-SAE versus not based on *blastN* alignments of the resulting sequence to an M121 reference. For 16 isolates without amplification products for sanger sequencing, we relied instead on whole genome sequencing to assess the presence or absence of the diagnostic HOU_2198 allele.

### Molecular clock analysis

A maximum likelihood phylogeny was constructed from a subset of all 155 USA300-SAE genomes with available dates and countries of sampling using the same method as above. The tree was rooted at HUV01 based on its position in the full ST8 phylogeny. *BactDating 1.1.2* was run on the rooted phylogeny using a fixed root and a strict gamma clock model to 1e8 MCMC iterations, sampled every 100,000 iterations (63). Effective sample sizes were evaluated with a 50% burn-in. Model fit and conversion were analyzed and compared to other models using the deviance information criterion and effective sample sizes as calculated by *BactDating* and *coda*, and *FigTree 1.4.4* was used to format output files for visualization in R (64).

### Ancestral reconstruction

We performed an ancestral reconstruction of geographic locations on the output tree from the molecular clock analysis using *PastML 1.9.43* using the default Marginal Posterior Probabilities Approximation(MPPA) method and the F81 evolutionary model (65). We visualized nodes with >50% marginal posterior probability for one country.

### Estimations of COMER coverage

Based on *blastN* alignments of contigs to the CA12 (NZ_CP007672.1) COMER sequence, we identified regions of the mobile element covered with >99% identity in each USA300-SAE genome included in the molecular clock and ancestral reconstruction analysis. For genomes missing the start of COMER, trimmed reads were directly aligned to CA12’s SCCmec to COMER region (NZ_CP007672.1:33712…78207) using *bowtie2 2.5.3* and read depth per position was calculated using *samtools 1.19.1*. (66,67).

### Experimental Strains

The deletion mutant M121-Δ*copX-mco-copL* was generated in an M121 background using an FPR3757 template, extending from 113 bp upstream of COMER’s *copX* through the beginning of the initial coding region of *copL*, resulting in a mutant with a complete deletion of *copX* and *mco* and a premature stop codon disrupting the *copL* open reading frame (termed M121Δ*copX-mco*). This mutant was constructed by allelic exchange using the temperature sensitive plasmid pIMAY, which uses a secY antisense counterselection determinant (68,69). The upstream fragment was amplified with primer 1F (CCG<u>GAATT</u>CCATATGTTGTTTCATTAGAGG) and 1R (AAGGTTTTCGATTTATCTCC) and the downstream fragment was amplified with primers 2F (GATATTAGGATCATTACTAATA)(2F) and 2R (AAGGAAAAAA<u>GCGGCCGC</u>ATAGGGCCTAATTTCAACTG) with *S.aureus* FPR3757 genomic DNA as template and PrimeStar Max DNA polymerase (Takara). After subcloning into pCR2.1 (Invitrogen), the fragment was cloned into pIMAY into EcoRI and NotI restriction sites. The recombinant plasmid was transformed into chemically competent E.coli SA08B (Lucigen). The plasmid was isolated from *E.coli*, verified by sequencing and then electroporated into *S.aureus* M121, which was plated onto TSA plus Cm 10 ug/mL and incubated at 28 °C. Transformants were incubated on TSA plus Cm 10 ug/mL at 37 °C to stimulate pIMAY chromosomal integration. Colonies free of replicating plasmid were grown at 28°C without Cm and then plated onto TSA containing anhydrotetracycline 1μg/mL for counterselection. Resulting colonies that showed loss of Cm resistance were screened by colony PCR for the absence of *copX*. The selected mutant was sequenced by Illumina whole genome sequencing (Plasmidsaurus) and demonstrated a complete deletion of *copX* and *mco* and a frameshift deletion causing a premature stop codon at AA17 of *copL*. The full *copX-mco-copL* sequence, including its native promoter, was complemented back into the mutant using the *erm* shuttle plasmid pCB07. pCB07 is pJC1306, which integrates into the SAPI1 attC site, with ermC replacing tetM (31).pCB07 was made using Gibson Assembly, where all of pJC1306 excluding tetM was amplified then assembled with *ermC* cassette amplified from pAH09(70). Starting 407 nt upstream of the start codon of copX on COMER and ending 458 nt downstream of the copL stop codon on COMER was amplified from M121 then put into BamHI-digested pCB07 using Gibson assembly to generate pCB07-COMER. pCB07 and pCB07-COMER were sequenced by Genewiz. Plasmids were electroporated into *S. aureus* RN4220 with the plasmid pRN7023 (strain RN9011), confirmed via PCR using primers JCO717 and JCO719, then transduced into M121-Δ*copX-mco-copL* via bacteriophage Φ80α.(31) Final constructs were sequenced by Plasmidsaurus.

### *In vitro* copper survival in water

Survival in aqueous copper was assayed in M121, M121Δ*copX-mco-copL*, the complemented M121Δ*copX-mco-copL::copX-mco-copL,* and M121Δ*copX-mco-copL* with an empty SAPI1 attC vector. Survival was measured by growing single colonies overnight in LB, subculturing overnight cultures to exponential phase, then normalizing each culture to 1.0 OD. Cultures were added to a 96-well plate, in technical duplicate, in a 1:1 ratio with a CuCl_2_ solution in molecular grade water to CuCl_2_ concentrations of 0 and 10 µM. Plates were incubated at room temperature for 24 hours, then spot plated in triplicate on LB and grown for 24 hours at 37°C. Results for the mutant were compared to the M121 background strain at each tested concentration using a pairwise Wilcoxon Mann-Whitney U test with a Benjamini-Hochberg correction for multiple comparisons.

### *In vitro* copper growth in RPMI-AA media

M121, the deletion mutant M121Δ*copX-mco-copL*, the complemented M121Δ*copX-mco-copL: copX-mco-copL,* and M121Δ*copX-mco-copL* with an empty shuttle vector transduced in place of the complementing construct were plated onto Luria broth agar (LBA) and grown overnight at 37°C. Single colonies were grown statically overnight at 37°C with 5% CO2 in RPMI 1640 tissue culture medium (Gibco 11835-030) supplemented with 2%(v/v) RPMI 1640 amino acid solution 50X (Sigma R7131). Growth assays were performed on overnight cultures in a round bottom 96-well plate. A 1M Copper (II) Chloride stock was made fresh for each experiment from Copper (II) Chloride powder (Sigma 203149-10G) and nuclease-free, deionized water (Growcells-GROW-NUPW-0500). The stock was used to make 4mM, 2mM, 1mM, 0.5mM, 0.25mM and 0.125mM in the RPMI 1640 + amino acid cultures at OD_600_ 0.025 in a final volume of 200µL per well. These subcultures were grown statically for 24H at 37°C with 5% CO2. Growth and blank media controls were run in the same assay plates. At 24 hours, plates were centrifuged at 4°C, 4000 RPM for 10 minutes. The supernatant was removed and the pellets were washed with nuclease free, deionized water. This process was repeated for a total of 2 washes to remove any traces of copper. Pellets were resuspended in 200uL of nuclease free, deionized water, then serially diluted and plated on LBA for CFU counting after overnight growth at 37°C.

## Data Visualization

The tree was visualized in ITOL (71). All other visualizations were made in *R 3.4.2* using *ape 5.7.1, ggtree 3.10.0*, *treeio 1.26.0, dplyr 1.1.4, ggplot2 3.5.0, phytools 2.3.0, ggnewscale 0.4.10, ggtreeExtra 1.12.0, ggstar 1.0.4, tidyr 1.3.1, reshape2 1.4.4, RColorBrewer 1.1.3, and stringr 1.5.1* and finished in Adobe Illustrator (72–75).

## Data & Code Availability

The scripts necessary to reproduce figures are available at https://github.com/amyecampbell/ChileUSA300. The raw reads generated for this project are available under NCBI BioProject Accession PRJNA1491352.

## Supporting information

Isolate Information

## ACKNOWLEDGEMENTS

The authors would like to acknowledge the Microbiome Sequencing Center, the Microbial Archive and Cryocollection (MicrobialARC), and the Center for Microbial Medicine, and the Research Institute at the Children’s Hospital of Philadelphia for sequencing support. This study was supported by the Agencia Nacional de Investigación y Desarrollo -ANID SENTINET CIN250062 (J.M.M. and L.D.), the US Centers for Disease Control and Prevention, Cooperative Agreement #6 NU3HCK000012-05-01 to J.M.M. (ARCH initiative), through Fondo Nacional de Desarrollo Científico y Tecnológico regular #1250588 to J.M.M. and #1260281 to L.D., and Fondo Nacional de Desarrollo Científico y Tecnológico Iniciación #11260350 to J.R.W.M. A.E.C. was supported by National Institutes of Health #K12GM081259.

---

**Figure S1.**
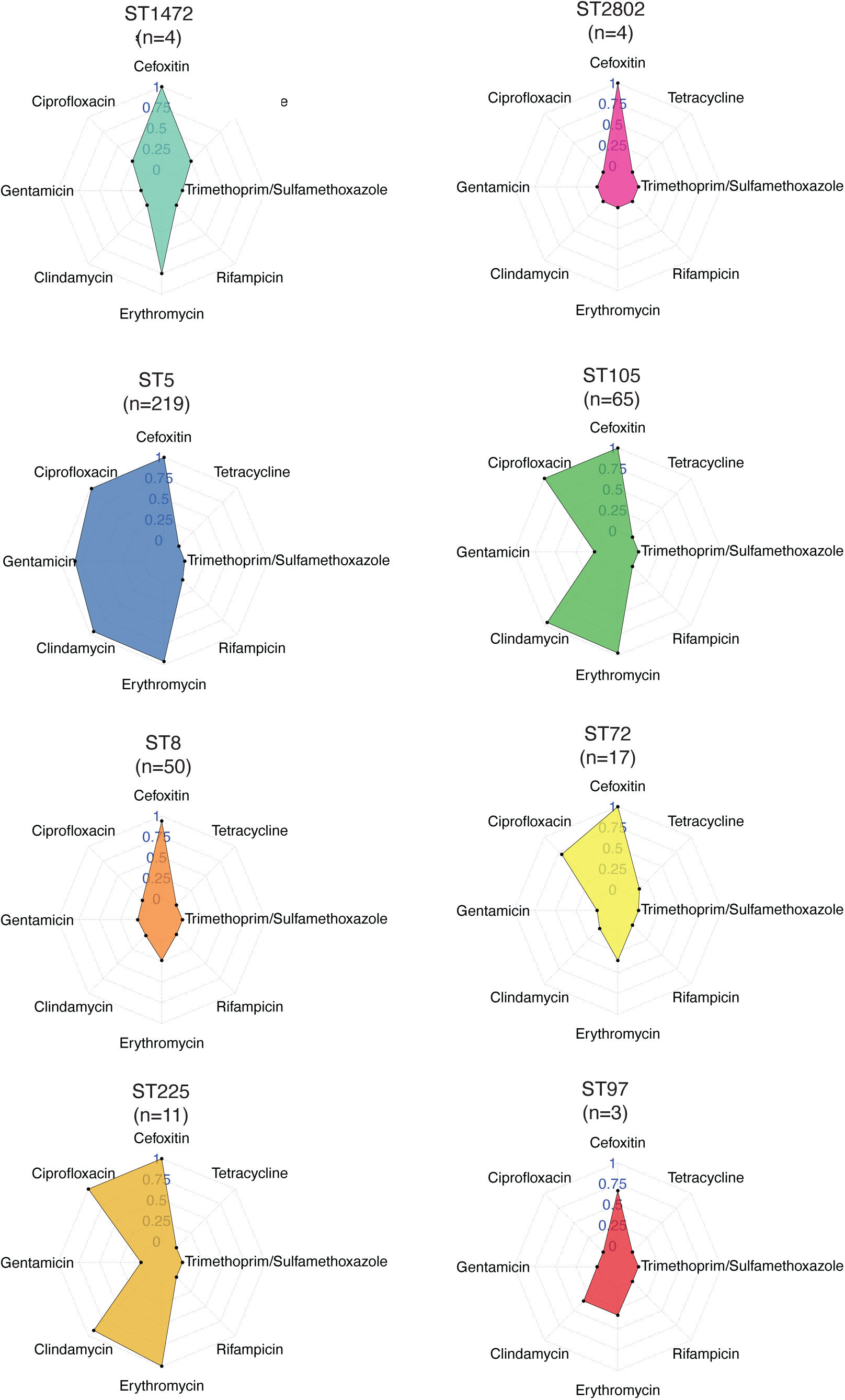
% of isolates in our collection resistant to each of 8 tested antibiotics.

**Figure S2.**
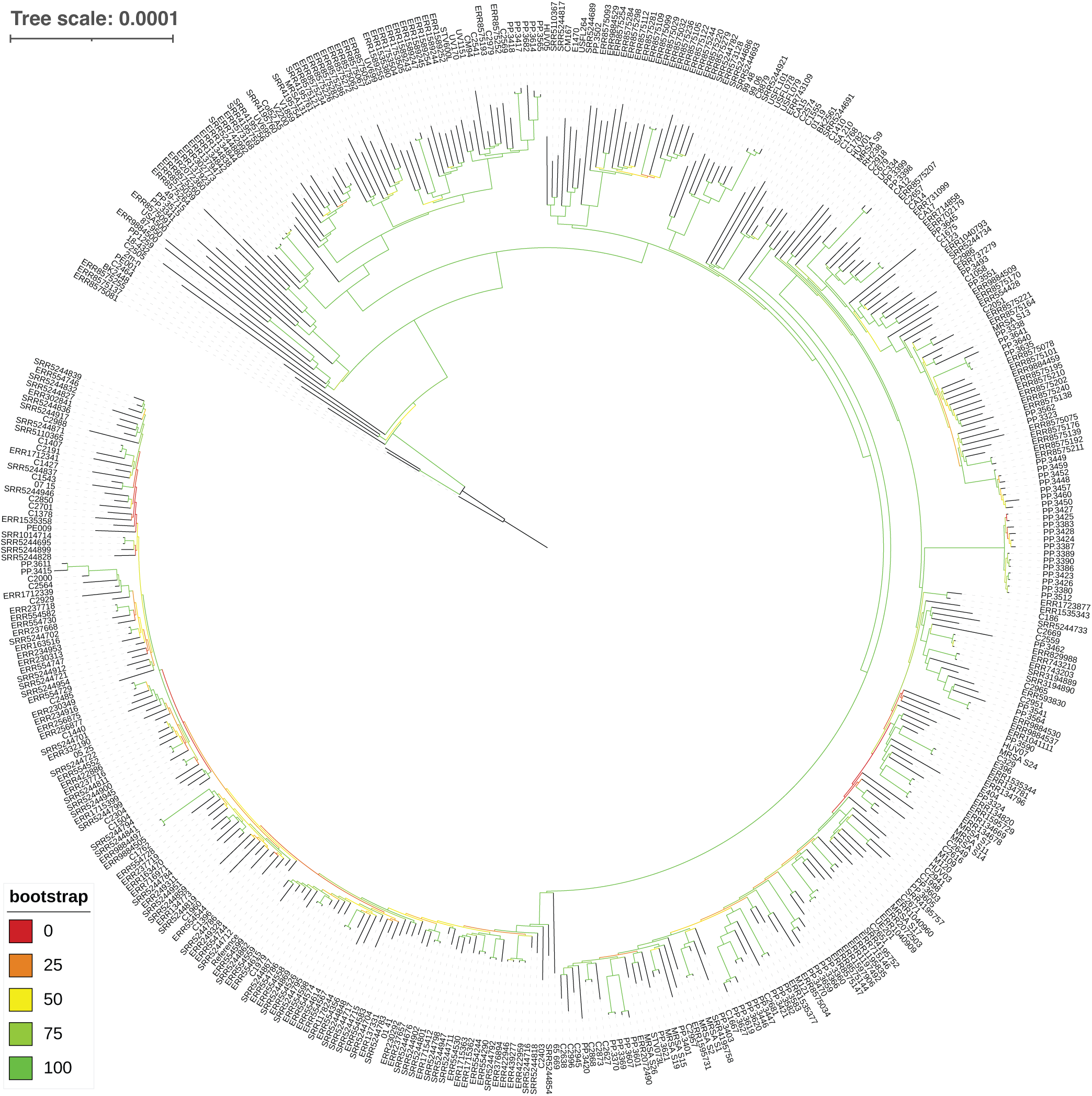
Related to. figure 2**. Bootstrap support for ST8 tree out of 100 bootstrap pseudoreplicates.**

**Figure S3.**
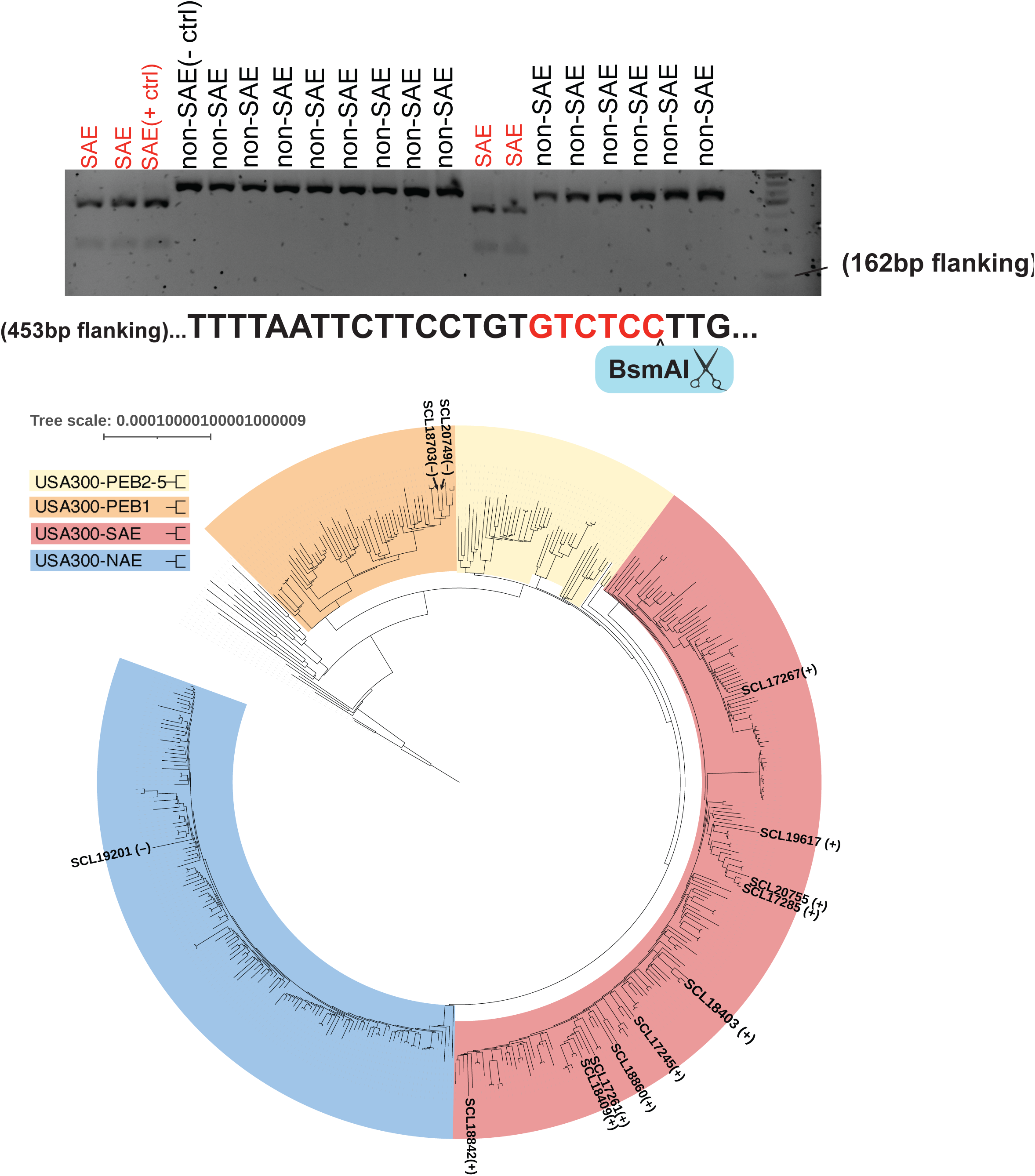
a) Schematic of CURED-based restriction digest method for identifying USA300-SAE among MRSA b) Maximum likelihood phylogeny showing phylogenetic placement of 13 ST8 clade isolates (parentheses indicate whether CURED restriction digest classified the isolates as SAE (+) or non-SAE (–)

**Figure S4.**
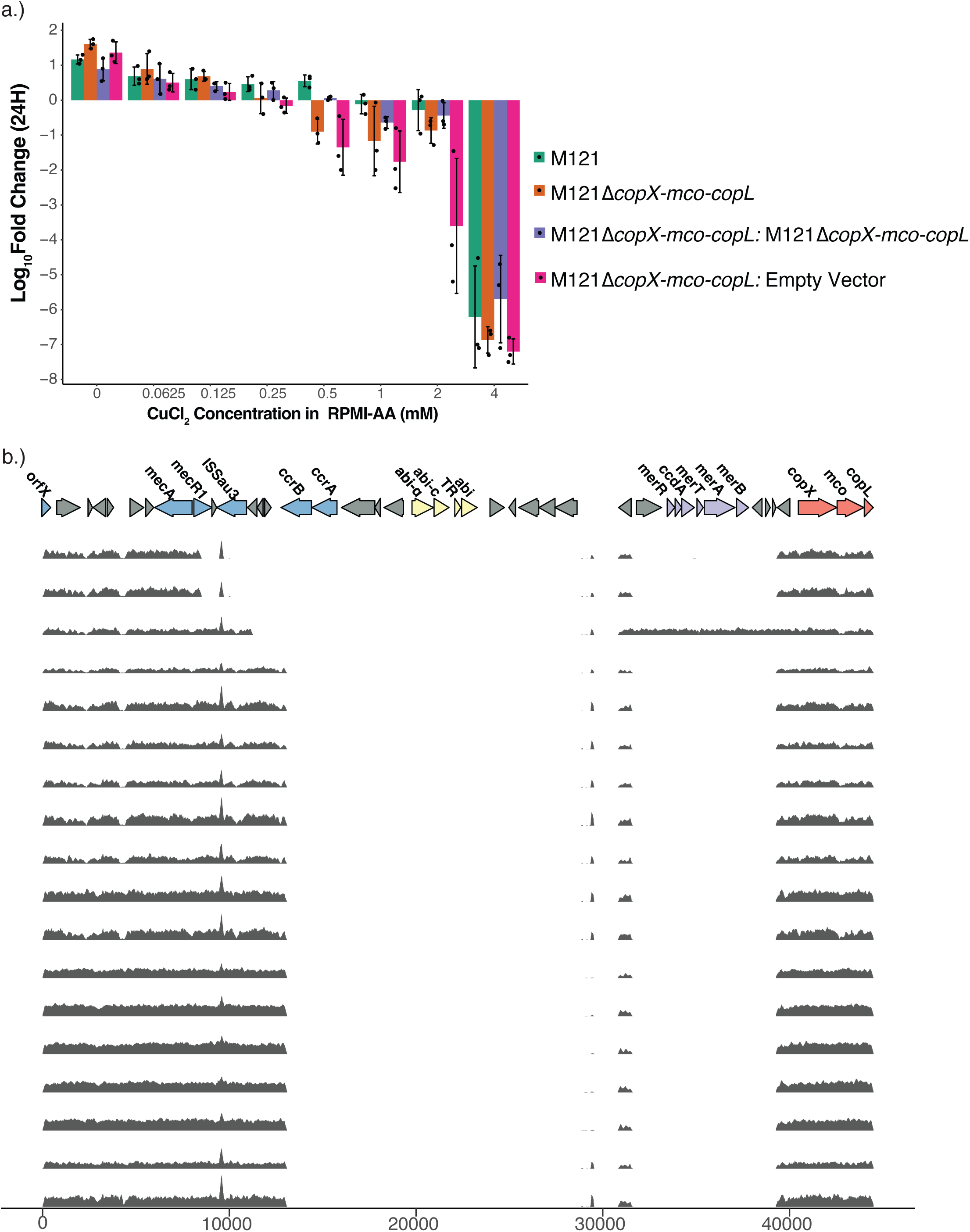
a) Growth in RPMI-AA media with copper at a range of concentrations. b) Relative depth of reads aligned to the COMER region among genomes missing the start of COMER

## REFERENCES

1. Parsons JB, Mourad A, Conlon BP, Kielian T, Fowler VG. Methicillin-resistant and susceptible Staphylococcus aureus: tolerance, immune evasion and treatment. Nat Rev Microbiol. 2025 Aug 20;1–19. doi:10.1038/s41579-025-01226-2

2. Chambers HF, DeLeo FR. Waves of resistance: Staphylococcus aureus in the antibiotic era. Nat Rev Microbiol. 2009 Sep;7(9):629–41. doi:10.1038/nrmicro2200

3. Arias CA, Reyes J, Carvajal LP, Rincon S, Diaz L, Panesso D, et al. A Prospective Cohort Multicenter Study of Molecular Epidemiology and Phylogenomics of Staphylococcus aureus Bacteremia in Nine Latin American Countries. Antimicrobial Agents and Chemotherapy. 2017 Sep 22;61(10):10.1128/aac.00816-17. doi:10.1128/aac.00816-17

4. Rodríguez-Noriega E, Seas C. The changing pattern of methicillin-resistant staphylococcus aureus clones in Latin America: implications for clinical practice in the region. Braz J Infect Dis. 2010 Dec;14:87–96. doi:10.1590/S1413-86702010000800004

5. Egea AL, Gagetti P, Lamberghini R, Faccone D, Lucero C, Vindel A, et al. New patterns of methicillin-resistant *Staphylococcus aureus* (MRSA) clones, community-associated MRSA genotypes behave like healthcare-associated MRSA genotypes within hospitals, Argentina. International Journal of Medical Microbiology. 2014 Nov 1;304(8):1086–99. doi:10.1016/j.ijmm.2014.08.002

6. Arias CA, Rincon S, Chowdhury S, Martínez E, Coronell W, Reyes J, et al. MRSA USA300 Clone and VREF — A U.S.–Colombian Connection? New England Journal of Medicine. 2008 Nov 13;359(20):2177–9. doi:10.1056/NEJMc0804021

7. Alvarez CA, Barrientes OJ, Leal AL, Contreras GA, Barrero L, Rincón S, et al. Community-associated Methicillin-resistant Staphylococcus aureus, Colombia - Volume 12, Number 12—December 2006 - Emerging Infectious Diseases journal - CDC [Internet]. doi:10.3201/eid1212.060814

8. Planet PJ, Diaz L, Kolokotronis SO, Narechania A, Reyes J, Xing G, et al. Parallel Epidemics of Community-Associated Methicillin-Resistant Staphylococcus aureus USA300 Infection in North and South America. The Journal of Infectious Diseases. 2015 Dec 15;212(12):1874–82. doi:10.1093/infdis/jiv320

9. Bianco CM, Moustafa AM, O’Brien K, Martin MA, Read TD, Kreiswirth BN, et al. Pre-epidemic evolution of the MRSA USA300 clade and a molecular key for classification. Frontiers in Cellular and Infection Microbiology. 2023;13. doi:10.3389/fcimb.2023.1081070

10. Aires de Sousa M, Miragaia M, Santos Sanches I, Ávila S, Adamson I, Casagrande ST, et al. Three-Year Assessment of Methicillin-ResistantStaphylococcus aureus Clones in Latin America from 1996 to 1998. Journal of Clinical Microbiology. 2001 Jun;39(6):2197–205. doi:10.1128/jcm.39.6.2197-2205.2001

11. Reyes J, Rincón S, Díaz L, Panesso D, Contreras GA, Zurita J, et al. Dissemination of Methicillin-Resistant Staphylococcus aureus USA300 Sequence Type 8 Lineage in Latin America. Clinical Infectious Diseases. 2009 Dec 15;49(12):1861–77. doi:10.1086/648426

12. Cabrejos-Hirashima L, Vives-Kufoy C, Inga-Salazar J, Astocondor L, Hinostroza N, García C. Frecuencia de Staphylococcus aureus meticilinorresistente adquirido en la comunidad en un hospital de tercer nivel en Perú. Rev Peru Med Exp Salud Publica. 2021 Jul 7;38(2):313–7. doi:10.17843/rpmesp.2021.382.6867

13. Martínez JRW, Planet PJ, Spencer-Sandino M, Rivas L, Díaz L, Moustafa AM, et al. Dynamics of the MRSA Population in a Chilean Hospital: a Phylogenomic Analysis (2000–2016). Microbiology Spectrum. 2023 Jun 20;11(4):e05351–22. doi:10.1128/spectrum.05351-22

14. Medina G, Egea AL, Otth C, Otth L, Fernández H, Bocco JL, et al. Molecular epidemiology of hospital-onset methicillin-resistant Staphylococcus aureus infections in Southern Chile. Eur J Clin Microbiol Infect Dis. 2013 Dec 1;32(12):1533–40. doi:10.1007/s10096-013-1907-8

15. Knight GM, Budd EL, Whitney L, Thornley A, Al-Ghusein H, Planche T, et al. Shift in dominant hospital-associated methicillin-resistant Staphylococcus aureus (HA-MRSA) clones over time. Journal of Antimicrobial Chemotherapy. 2012 Oct 1;67(10):2514–22. doi:10.1093/jac/dks245

16. Planet PJ, LaRussa SJ, Dana A, Smith H, Xu A, Chanelle Ryan, et al. Emergence of the Epidemic Methicillin-Resistant Staphylococcus aureus Strain USA300 Coincides with Horizontal Transfer of the Arginine Catabolic Mobile Element and speG-mediated Adaptations for Survival on Skin. mBio. 2013 Dec 17;4(6):10.1128/mbio.00889-13. doi:10.1128/mbio.00889-13

17. Thurlow LR, Joshi GS, Clark JR, Spontak JS, Neely CJ, Maile R, et al. Functional Modularity of the Arginine Catabolic Mobile Element Contributes to the Success of USA300 Methicillin-Resistant Staphylococcus aureus. Cell Host & Microbe. 2013 Jan 16;13(1):100–7. doi:10.1016/j.chom.2012.11.012 PubMed PMID: 23332159.

18. Baldan R, Rancoita PMV, Di Serio C, Mazzotti M, Cichero P, Ossi C, et al. Epidemic MRSA clone ST22-IV is more resistant to multiple host- and environment-related stresses compared with ST228-I. Journal of Antimicrobial Chemotherapy. 2015 Mar 1;70(3):757–65. doi:10.1093/jac/dku467

19. Copin R, Sause WE, Fulmer Y, Balasubramanian D, Dyzenhaus S, Ahmed JM, et al. Sequential evolution of virulence and resistance during clonal spread of community-acquired methicillin-resistant Staphylococcus aureus. PNAS. 2019 Jan 29;116(5):1745–54. doi:10.1073/pnas.1814265116 PubMed PMID: 30635416.

20. McDougal LK, Steward CD, Killgore GE, Chaitram JM, McAllister SK, Tenover FC. Pulsed-Field Gel Electrophoresis Typing of Oxacillin-Resistant *Staphylococcus aureus* Isolates from the United States: Establishing a National Database. Journal of Clinical Microbiology. 2003;41(11):5113–20. doi:10.1128/jcm.41.11.5113-5120.2003

21. Aanensen DM, Feil EJ, Holden MTG, Dordel J, Yeats CA, Fedosejev A, et al. Whole-Genome Sequencing for Routine Pathogen Surveillance in Public Health: a Population Snapshot of Invasive Staphylococcus aureus in Europe. mBio. 2016;7(3):10.1128/mbio.00444-16. doi:10.1128/mbio.00444-16

22. Di Gregorio S, Vielma J, Haim MS, Rago L, Campos J, Kekre M, et al. Genomic epidemiology of Staphylococcus aureus isolated from bloodstream infections in South America during 2019 supports regional surveillance. Microbial Genomics. 2023;9(5):001020. doi:10.1099/mgen.0.001020

23. Esteves MAC, Viana AS, Viçosa GN, Botelho AMN, Moustafa AM, Mansoldo FRP, et al. RdJ detection tests to identify a unique MRSA clone of ST105-SCCmecII lineage and its variants disseminated in the metropolitan region of Rio de Janeiro. Frontiers in Microbiology. 2023;14. doi:10.3389/fmicb.2023.1275918

24. Viana AS, Botelho AMN, Moustafa AM, Boge CLK, Ferreira ALP, Carvalho MC da S, et al. Multidrug-Resistant Methicillin-Resistant Staphylococcus aureus Associated with Bacteremia and Monocyte Evasion, Rio de Janeiro, Brazil - Volume 27, Number 11—November 2021 - Emerging Infectious Diseases journal - CDC [Internet]. doi:10.3201/eid2711.210097

25. Theiller E, Rajagopol S, Higgins S, Torres DI, Napper T, Galis BE, et al. Empowering global disease surveillance with CURED: a tool for rapid identification of unique genomic biomarkers. mSystems. 2026 Mar 19;11(4):e01063–25. doi:10.1128/msystems.01063-25

26. Aguayo-Reyes A, Morales-León F, Quezada-Aguiluz M, Opazo-Capurro A, Bello-Toledo H, Mella S, et al. Emergence of ST8-USA300 and ST8-USA300-Latin American variant: a changing landscape of community-associated methicillin-resistant Staphylococcus aureus in Chile. Microbiology Spectrum. 2025 Sep 30;13(11):e01031–25. doi:10.1128/spectrum.01031-25

27. Bletz S, Mellmann A, Rothgänger J, Harmsen D. Ensuring backwards compatibility: traditional genotyping efforts in the era of whole genome sequencing. Clinical Microbiology and Infection. 2015 Apr 1;21(4):347.e1-347.e4. doi:10.1016/j.cmi.2014.11.005 PubMed PMID: 25658529.

28. Zapotoczna M, Riboldi GP, Moustafa AM, Dickson E, Narechania A, Morrissey JA, et al. Mobile-Genetic-Element-Encoded Hypertolerance to Copper Protects Staphylococcus aureus from Killing by Host Phagocytes. mBio. 2018 Oct 16;9(5):10.1128/mbio.00550-18. doi:10.1128/mbio.00550-18

29. Rosario-Cruz Z, Eletsky A, Daigham NS, Al-Tameemi H, Swapna GVT, Kahn PC, et al. The *copBL* operon protects *Staphylococcus aureus* from copper toxicity: CopL is an extracellular membrane–associated copper-binding protein. Journal of Biological Chemistry. 2019 Mar 15;294(11):4027–44. doi:10.1074/jbc.RA118.004723

30. Purves J, Thomas J, Riboldi GP, Zapotoczna M, Tarrant E, Andrew PW, et al. A horizontally gene transferred copper resistance locus confers hyper-resistance to antibacterial copper toxicity and enables survival of community acquired methicillin resistant Staphylococcus aureus USA300 in macrophages. Environmental Microbiology. 2018;20(4):1576–89. doi:10.1111/1462-2920.14088

31. Chen J, Yoong P, Ram G, Torres VJ, Novick RP. Single-copy vectors for integration at the SaPI1 attachment site for *Staphylococcus aureus*. Plasmid. 2014 Nov 1;76:1–7. doi:10.1016/j.plasmid.2014.08.001

32. World Health Organization, editor. Guidelines for drinking-water quality. Fourth edition incorporating the first and second addenda. Geneva: World Health Organization; 2022. 1 p.

33. Planet PJ, Diaz L, Rios R, Arias CA. Global Spread of the Community-Associated Methicillin-Resistant Staphylococcus aureus USA300 Latin American Variant. The Journal of Infectious Diseases. 2016 Nov 15;214(10):1609–10. doi:10.1093/infdis/jiw418

34. Aguayo-Reyes A, Morales-León F, Quezada-Aguiluz M, Quezada M, Mella S, Riedel G, et al. Community-acquired methicillin-resistant Staphylococcus aureus invasive infections: a case series from Central-South Chile. Front Med. 2024 May 15;11. doi:10.3389/fmed.2024.1365756

35. Porse A, Schou TS, Munck C, Ellabaan MMH, Sommer MOA. Biochemical mechanisms determine the functional compatibility of heterologous genes. Nat Commun. 2018 Feb 6;9(1):522. doi:10.1038/s41467-018-02944-3

36. Rajer F, Sandegren L. The Role of Antibiotic Resistance Genes in the Fitness Cost of Multiresistance Plasmids. mBio. 2022 Jan 18;13(1):e03552–21. doi:10.1128/mbio.03552-21

37. Dorado-Morales P, Garcillán-Barcia MP, Lasa I, Solano C. Fitness Cost Evolution of Natural Plasmids of Staphylococcus aureus. mBio. 2021 Feb 23;12(1):10.1128/mbio.03094-20. doi:10.1128/mbio.03094-20

38. Harkins CP, Pichon B, Doumith M, Parkhill J, Westh H, Tomasz A, et al. Methicillin-resistant Staphylococcus aureus emerged long before the introduction of methicillin into clinical practice. Genome Biology. 2017 Jul 20;18(1):130. doi:10.1186/s13059-017-1252-9

39. Tapia J, Pía Rodríguez M, Castillo P, González R, Rodríguez C, Valdés A, et al. Chapter 7: Arsenic and Copper in Chile and the Development of Environmental Standards. In: Chile: Environmental Perspectives and Challenges. Hauppauge, NY: Nova Science Publishers, Inc.; 2019. (Environmental Research Advances).

40. Romero L, Alonso H, Campano P, Fanfani L, Cidu R, Dadea C, et al. Arsenic enrichment in waters and sediments of the Rio Loa (Second Region, Chile). Applied Geochemistry. 2003 Sep 1;Arsenic Geochemistry-selected papers from the 10th Water-Rock Interaction Symposium, Villasimius, Italy, 10-15 June 200118(9):1399–416. doi:10.1016/S0883-2927(03)00059-3

41. Jelinkova P, Vesely R, Cihalova K, Hegerova D, Ananbeh HAAA, Richtera L, et al. Effect of arsenic (III and V) on oxidative stress parameters in resistant and susceptible *Staphylococcus aureus*. Environmental Research. 2018 Oct 1;166:394–401. doi:10.1016/j.envres.2018.06.024

42. Baker J, Sitthisak S, Sengupta M, Johnson M, Jayaswal RK, Morrissey JA. Copper Stress Induces a Global Stress Response in Staphylococcus aureus and Represses sae and agr Expression and Biofilm Formation. Applied and Environmental Microbiology. 2010 Jan;76(1):150–60. doi:10.1128/AEM.02268-09

43. Martínez JRW, Alcalde-Rico M, Jara-Videla E, Reyes J, Carvajal LP, Rincon S, et al. Geographic divergence of methicillin-resistant Staphylococcus aureus ST5-SCCmecI in the aftermath of a major earthquake and tsunami: impact of a plasmid harboring heavy metal resistance genes. mBio. 2025 Mar 19;16(4):e03669–24. doi:10.1128/mbio.03669-24

44. Strauß L, Stegger M, Akpaka PE, Alabi A, Breurec S, Coombs G, et al. Origin, evolution, and global transmission of community-acquired Staphylococcus aureus ST8. Proceedings of the National Academy of Sciences. 2017 Dec 5;114(49):E10596–604. doi:10.1073/pnas.1702472114

45. Committee C in DW, Staff NRC, Staff C on LS. Copper in Drinking Water. Washington: National Academies Press; 2000. 162 p.

46. Saenkham-Huntsinger P, Hyre AN, Hanson BS, Donati GL, Adams LG, Ryan C, et al. Copper Resistance Promotes Fitness of Methicillin-Resistant Staphylococcus aureus during Urinary Tract Infection. mBio. 2021 Sep 7;12(5):10.1128/mbio.02038-21. doi:10.1128/mbio.02038-21

47. Argudín MA, Deplano A, Nonhoff C, Yin N, Michel C, Martiny D, et al. Epidemiology of the Staphylococcus aureus CA-MRSA USA300 in Belgium. Eur J Clin Microbiol Infect Dis. 2021 Nov 1;40(11):2335–47. doi:10.1007/s10096-021-04286-3

48. Sassi M, Felden B, Revest M, Tattevin P, Augagneur Y, Donnio PY. An outbreak in intravenous drug users due to USA300 Latin-American variant community-acquired methicillin-resistant Staphylococcus aureus in France as early as 2007. Eur J Clin Microbiol Infect Dis. 2017 Dec 1;36(12):2495–501. doi:10.1007/s10096-017-3092-7

49. Nurjadi D, Fleck R, Lindner A, Schäfer J, Gertler M, Mueller A, et al. Import of community-associated, methicillin-resistant *Staphylococcus aureus* to Europe through skin and soft-tissue infection in intercontinental travellers, 2011–2016. Clinical Microbiology and Infection. 2019 Jun 1;25(6):739–46. doi:10.1016/j.cmi.2018.09.023

50. Nandi A, Pecetta S, Bloom DE. Global antibiotic use during the COVID-19 pandemic: analysis of pharmaceutical sales data from 71 countries, 2020–2022. eClinicalMedicine. 2023 Mar 1;57. doi:10.1016/j.eclinm.2023.101848 PubMed PMID: 36776504.

51. Klein EY, Impalli I, Poleon S, Denoel P, Cipriano M, Van Boeckel TP, et al. Global trends in antibiotic consumption during 2016–2023 and future projections through 2030. Proceedings of the National Academy of Sciences. 2024 Dec 3;121(49):e2411919121. doi:10.1073/pnas.2411919121

52. Clarke EL, Taylor LJ, Zhao C, Connell A, Lee JJ, Fett B, et al. Sunbeam: an extensible pipeline for analyzing metagenomic sequencing experiments. Microbiome. 2019 Mar 22;7(1):46. doi:10.1186/s40168-019-0658-x

53. Krueger F, James F, Ewels P, Afyounian E, Weinstein M, Schuster-Boeckler B, et al. TrimGalore: v0.6.10 [Internet]. 2023 [cited 2024 Oct 10]. Available from: https://zenodo.org/records/7598955 doi:10.5281/zenodo.7598955

54. Seemann T. Shovill [Internet]. Available from: https://github.com/tseemann/shovill

55. Parks DH, Imelfort M, Skennerton CT, Hugenholtz P, Tyson GW. CheckM: assessing the quality of microbial genomes recovered from isolates, single cells, and metagenomes. Genome Res. 2015 Jul 1;25(7):1043–55. doi:10.1101/gr.186072.114 PubMed PMID: 25977477.

56. Ondov BD, Starrett GJ, Sappington A, Kostic A, Koren S, Buck CB, et al. Mash Screen: high-throughput sequence containment estimation for genome discovery. Genome Biology. 2019 Nov 5;20(1):232. doi:10.1186/s13059-019-1841-x

57. Jolley KA, Maiden MC. BIGSdb: Scalable analysis of bacterial genome variation at the population level. BMC Bioinformatics. 2010 Dec 10;11(1):595. doi:10.1186/1471-2105-11-595

58. Seemann T. mlst [Internet]. Available from: https://github.com/tseemann/mlst

59. Stamatakis A. RAxML version 8: a tool for phylogenetic analysis and post-analysis of large phylogenies. Bioinformatics. 2014 May 1;30(9):1312–3. doi:10.1093/bioinformatics/btu033

60. Didelot X, Wilson DJ. ClonalFrameML: Efficient Inference of Recombination in Whole Bacterial Genomes. PLOS Computational Biology. 2015 Feb 12;11(2):e1004041. doi:10.1371/journal.pcbi.1004041

61. Schwengers O, Jelonek L, Dieckmann MA, Beyvers S, Blom J, Goesmann A. Bakta: rapid and standardized annotation of bacterial genomes via alignment-free sequence identification. Microbial Genomics. 2021;7(11):000685. doi:10.1099/mgen.0.000685

62. O’Leary NA, Cox E, Holmes JB, Anderson WR, Falk R, Hem V, et al. Exploring and retrieving sequence and metadata for species across the tree of life with NCBI Datasets. Sci Data. 2024 Jul 5;11(1):732. doi:10.1038/s41597-024-03571-y

63. Didelot X, Croucher NJ, Bentley SD, Harris SR, Wilson DJ. Bayesian inference of ancestral dates on bacterial phylogenetic trees. Nucleic Acids Research. 2018 Dec 14;46(22):e134. doi:10.1093/nar/gky783

64. Plummer M, Best N, Cowles K, Vines K. coda: Output Analysis and Diagnostics for MCMC. 2005.

65. Ishikawa SA, Zhukova A, Iwasaki W, Gascuel O. A Fast Likelihood Method to Reconstruct and Visualize Ancestral Scenarios. Molecular Biology and Evolution. 2019 Sep 1;36(9):2069–85. doi:10.1093/molbev/msz131

66. Langmead B, Salzberg SL. Fast gapped-read alignment with Bowtie 2. Nat Methods. 2012 Apr;9(4):4. doi:10.1038/nmeth.1923

67. Li H, Handsaker B, Wysoker A, Fennell T, Ruan J, Homer N, et al. The Sequence Alignment/Map format and SAMtools. Bioinformatics. 2009 Aug 15;25(16):2078–9. doi:10.1093/bioinformatics/btp352

68. Monk IR, Foster TJ. Genetic manipulation of Staphylococci – Breaking through the barrier. Front Cell Infect Microbiol. 2012 Apr 12;2. doi:10.3389/fcimb.2012.00049

69. Monk IR, Shah IM, Xu M, Tan MW, Foster TJ. Transforming the Untransformable: Application of Direct Transformation To Manipulate Genetically Staphylococcus aureus and Staphylococcus epidermidis. mBio. 2012 Mar 20;3(2):10.1128/mbio.00277-11. doi:10.1128/mbio.00277-11

70. Boles BR, Horswill AR. agr-Mediated Dispersal of Staphylococcus aureus Biofilms. PLOS Pathogens. 2008 Apr 25;4(4):e1000052. doi:10.1371/journal.ppat.1000052

71. Letunic I, Bork P. Interactive Tree Of Life (iTOL) v5: an online tool for phylogenetic tree display and annotation. Nucleic Acids Research. 2021 Jul 2;49(W1):W293–6. doi:10.1093/nar/gkab301

72. Yu G. Using ggtree to Visualize Data on Tree-Like Structures. Current Protocols in Bioinformatics. 2020;69(1):e96. doi:10.1002/cpbi.96

73. Wang LG, Lam TTY, Xu S, Dai Z, Zhou L, Feng T, et al. Treeio: An R Package for Phylogenetic Tree Input and Output with Richly Annotated and Associated Data. Molecular Biology and Evolution. 2020 Feb 1;37(2):599–603. doi:10.1093/molbev/msz240

74. Paradis E, Schliep K. ape 5.0: an environment for modern phylogenetics and evolutionary analyses in R. Bioinformatics. 2019 Feb 1;35(3):526–8. doi:10.1093/bioinformatics/bty633

75. Revell LJ. phytools 2.0: an updated R ecosystem for phylogenetic comparative methods (and other things). PeerJ. 2024 Jan 5;12:e16505. doi:10.7717/peerj.16505

